# Subspace Image Synthesis for Style-Invariant Object Recognition

**DOI:** 10.64898/2026.10.21.759598

**Authors:** Hayato Ono, Ayumu Yamashita, Kaoru Amano, Masataka Sawayama

## Abstract

Human-like behavior in visual models does not necessarily imply human-like internal computation. When a model recognizes objects robustly across changes in visual style, what internal computations support this behavior, and which are relevant to human perception? We addressed this question by pruning style-robust neural networks into candidate submodels corresponding to two computational strategies. The first retained channels whose responses were stable across style changes for fixed content, yielding a Content Stability Score (CSS)-based submodel. The second retained channels whose responses were stable across content changes for fixed style, yielding a Style Stability Score (SSS)-based submodel. Across ImageNet, Stylized ImageNet, and model-versus-human benchmarks, the two submodel families showed comparable object-recognition performance. We then used their exclusive feature subspaces to generate matched images with Gram-matrix-based neural style transfer and asked human observers which image was more similar in visual style to a reference image. Images generated from the CSS-based submodel were selected significantly more often than chance, even though CSS preferentially retained channels whose individual activation patterns were relatively insensitive to style. These results suggest that representations preserving content across style changes are more consistent with human judgments of visual style. More broadly, they demonstrate a framework for isolating candidate computational subspaces from large artificial models and testing their perceptual relevance through image synthesis.

## Introduction

Machine learning models increasingly exhibit human-like behavior across a variety of visual tasks. However, similarity in behavior does not necessarily imply similarity in underlying computation. Modern artificial intelligence models contain many interacting representations and computational pathways, any subset of which may contribute to successful task performance. As a result, a model may achieve human-like behavior through a combination of mechanisms, some potentially shared with human vision and others entirely artificial. This raises the fundamental question of which internal computational functions underlie a model’s apparent alignment with human perception.

In this study, we propose a framework for addressing this question and apply it to investigate the computational principles underlying style-invariant object recognition (Geirhos et al., 2019, 2021). Specifically, we investigate computational subspaces within deep artificial neural networks through image synthesis. Below, we first describe the central idea of this framework and then explain how it can be applied to style-invariant object recognition.

Our approach builds on the literature of metamer research (Freeman & Simoncelli, 2011; Freeman, Ziemba, Heeger, Simoncelli, & Movshon, 2013; Wallis et al., 2019; Jagadeesh & Gardner, 2022; Feather, Leclerc, Mądry, & McDermott, 2023; Kamao, Ono, Yamashita, Amano, & Sawayama, 2026). In metamer studies, researchers assume an internal model of visual processing and synthesize stimuli that produce equivalent internal representations. The synthesized stimuli are then compared with real inputs, and the model is iteratively refined to better explain perceptual equivalence. For example, Freeman et al. revealed representational properties of peripheral vision by synthesizing images that matched V2-like feature representations (Freeman & Simoncelli, 2011; Freeman et al., 2013). More recently, neural networks have been used as models of internal visual processing to investigate whether synthesized metamers produce neural responses equivalent to those evoked by natural images in higher visual areas (Jagadeesh & Gardner, 2022).

Like metamer research, our approach evaluates alignment with human perception by asking how closely images synthesized using a candidate model resemble their target images perceptually. However, rather than treating a large pretrained neural network as a single fixed model, we examine different computational subspaces within the network. Specifically, we propose using pruning to construct multiple candidate submodels. We then synthesize images using each submodel and assess which submodels impose constraints that better capture human perception. Comparing the perceptual consequences of these submodel-specific constraints is the central idea of our approach.

We apply this framework to style-invariant object recognition (Geirhos et al., 2019, 2021). Humans can reliably recognize objects across substantial variations in visual style, including changes in material appearance, texture, illumination, and artistic rendering. This ability is essential because objects in natural environments are encountered under diverse visual appearances. However, classic neural network models often failed to recognize objects when their visual style changes. Instead of using shape information as humans typically do, they relied heavily on texture cues, a phenomenon known as texture bias. To address this limitation, numerous training strategies have been proposed, including style augmentation (Geirhos et al., 2019), adversarial training (Engstrom et al., 2019), and large-scale pretraining (Raghu, Unterthiner, Kornblith, Zhang, & Dosovitskiy, 2021; Radford et al., 2021). As a result, modern vision models exhibit substantially improved robustness to style variations. Nevertheless, it remains unclear what computational mechanisms within these robust models contribute to style-invariant recognition and which of those mechanisms are most consistent with human perception.

Our framework addresses this question by constructing submodels based on two candidate strategies for style-invariant object recognition. Each strategy defines a different criterion for pruning the same pretrained model. Specifically, we use activation-based structured pruning (Cheng, Zhang, & Shi, 2024), in which channels are retained or removed according to their response properties measured using a controlled dataset designed for each strategy. This procedure allows us to construct submodels that preferentially retain channels associated with different candidate representational functions.

The first strategy emphasizes channels whose responses remain stable across changes in visual style. To identify such channels, we constructed a dataset in which the same object content was rendered in multiple styles and defined a Content Stability Score (CSS) that quantified the stability of each channel’s activation pattern across these style changes. Channels with high CSS may encode object content or geometric structure, and are analogous to neurons in higher visual areas that preserve selectivity for object identity despite substantial variations in appearance (Srinath et al., 2021). This strategy reflects the idea that robust recognition can be achieved through representations that are largely invariant to style.

The second strategy emphasizes channels whose responses remain stable across changes in object content. Rather than eliminating style information, a recognition system may estimate style-related variables and use them to contextualize or normalize object representations. To identify such channels, we constructed a complementary dataset in which the same visual style was applied to multiple object contents and defined a Style Stability Score (SSS) that quantified the stability of each channel’s activation pattern across these content changes. Channels with high SSS may encode style-related information that remains consistent across different objects rendered in the same style. Consistent with this possibility, neurophysiological studies have identified neurons selective for material properties such as gloss, suggesting that the visual system may contain representations of style-related attributes (Nishio, Goda, & Komatsu, 2012).

Based on these possible computational strategies, we constructed submodels through activation-based structured pruning and synthesized stimuli from the resulting computational subspaces (Figure 1). In the following sections, we first describe how the submodels were constructed and evaluate how pruning affects object recognition performance relative to the original model (Experiment 1). We then conduct a psychophysical experiment to assess which submodel generates images that more closely align with human perception (Experiment 2). Through these experiments, we aim to identify computational principles that better account for human style-invariant object recognition.

**Figure 1:**
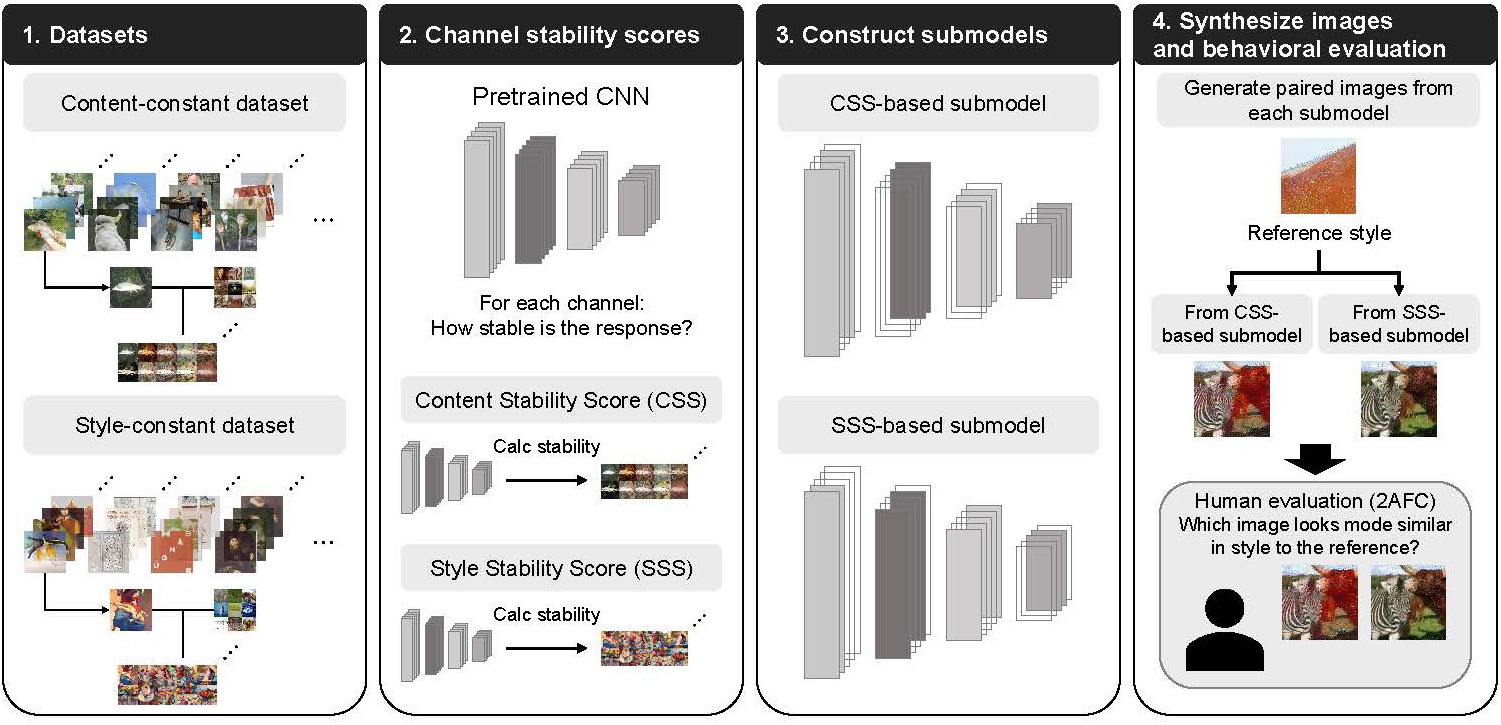
(1) Two controlled datasets were constructed to isolate complementary response properties: a content-constant dataset, in which object content was held fixed while visual style varied, and a style-constant dataset, in which visual style was held fixed while object content varied. (2) Channel-wise response stability was measured in convolutional neural networks using these datasets, yielding the Content Stability Score (CSS) and Style Stability Score (SSS), respectively. (3) Channels were ranked using the corresponding stability scores, and score-based structured pruning was used to construct CSS-based and SSS-based submodels that preferentially retained channels associated with each candidate computational strategy. (4) Matched images were synthesized from the two submodel-derived feature subspaces, and human observers performed a two-alternative forced-choice (2AFC) task to judge which synthesized image was more similar in visual style to the reference style image. All images shown in this figure were either obtained from ImageNet or WikiArt, or generated by applying style transfer to images from these sources.

## Experiment 1: Construction and evaluation of computational submodels

We first constructed submodels based on two candidate computational strategies. Before using these submodels to synthesize stimuli for the behavioral experiment, we evaluated how these models differed in object recognition performance. This section begins by describing the base models and then explains how the submodels were constructed. We then report evaluations of these submodels using several object-recognition and model–human comparison metrics.

### Base models: SIN-trained neural networks

Following Geirhos et al. (2019), we used ImageNet-pretrained neural networks fine-tuned on Stylized ImageNet as the base models for submodel construction to ensure meaningful object-recognition performance under style variation. Although some pre-trained models are shared by the authors, we trained our own Stylized ImageNet models because the present study required comparable feature models across multiple architectures. Stylized ImageNet was constructed by applying style transfer to ImageNet training data, which contains approximately 1.28 million images, using 79,395 WikiArt images as style references. For each ImageNet image, a WikiArt style image was randomly sampled, and a stylized image was generated using Adaptive Instance Normalization (AdaIN)-based style transfer (Huang & Belongie, 2017). The original ImageNet class labels were retained for supervised training.

We evaluated four architectures: AlexNet (Krizhevsky, Sutskever, & Hinton, 2017), VGG-19 (Simonyan & Zisserman, 2015), ResNet-50 (He, Zhang, Ren, & Sun, 2016), and CORnet-S (Kubilius et al., 2019). Each model was initialized from an ImageNet-pretrained checkpoint and then fine-tuned on the stylized training images using the original ImageNet labels. Fine-tuning was performed for 50 epochs using the SGD optimizer with a learning rate of 0.001, a momentum of 0.9, a batch size of 1024, and a weight decay of 0.0001. Model checkpoints were selected based on top-1 classification accuracy on the internal Stylized ImageNet validation set derived from the stylized ImageNet training split. The held-out stylized images (50,000 images) derived from the original ImageNet validation split were used only for final model-performance evaluation.

### Submodel construction

#### Datasets for constructing two submodels

We constructed submodels according to two broad strategies. The first strategy was to retain channels whose responses remained stable across changes in visual style. The second strategy was to retain channels whose responses remained stable across changes in object content. Here, we first describe the controlled datasets used to compute response-stability scores for these two strategies. One dataset consisted of image groups in which object content was held constant while visual style varied (Figure 2 (A)). The other consisted of image groups in which visual style was held constant while object content varied (Figure 2 (B)). We refer to the former as the content-constant dataset and the latter as the style-constant dataset.

**Figure 2:**
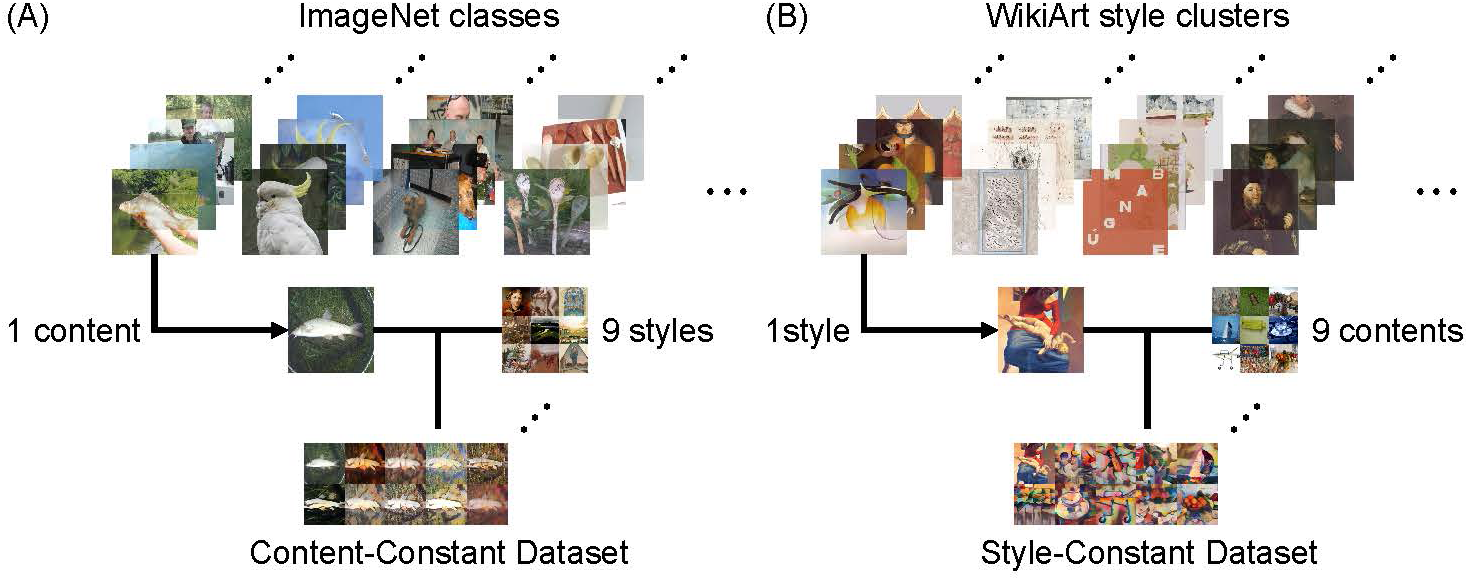
Overview of the content-constant and style-constant datasets. (A) In the content-constant dataset, one content image was selected from each of 1,000 ImageNet categories and stylized using 9 different WikiArt style images. Each group therefore consisted of 1 original content image and 9 stylized versions that preserved the same object content while varying in visual style. (B) In the style-constant dataset, 1,000 representative style images were selected from WikiArt by clustering VGG-16 feature representations and choosing the image closest to each cluster centroid. Each representative style image was applied to 9 different ImageNet content images. Each group therefore consisted of 1 original style image and 9 stylized images that shared the same visual style while varying in object content. Each dataset comprised 1,000 groups of 10 images—one original reference image and nine stylized images—yielding 10,000 images in total, including 9,000 generated stylized images. All images shown in this figure were either obtained from ImageNet or WikiArt, or generated by applying style transfer to images from these sources.

The two datasets were matched in the number of groups and in the number of stylized images per group. The content-constant dataset was based on 1,000 ImageNet object categories. For each category, one content image was selected and stylized with nine different WikiArt style images, yielding 1,000 original ImageNet images and 9,000 stylized images in total. Conversely, the style-constant dataset was based on 1,000 representative WikiArt style images. To sample these representative style images, we used K-means clustering. Specifically, we extracted activation patterns from the relu4_1 layer of a VGG-16 network (Simonyan & Zisserman, 2015) for all 79,395 WikiArt images and clustered these feature representations into 1,000 clusters. The image closest to each cluster centroid in this feature space was selected as the representative style image. For each representative style image, nine ImageNet content images were randomly sampled from the ImageNet training set and stylized with the same style image using AdaIN. This procedure yielded 1,000 original WikiArt images and 9,000 stylized images in total, matching the size of the content-constant dataset.

#### Activation-based pruning scores: Content Stability Score and Style Stability Score

Next, we describe how channel-wise activation patterns were converted into response-stability scores, Content Stability Score and Style Stability Score. For each dataset, we computed the corresponding stability score in two steps. In the first step, we quantified how stable each channel response was within each stimulus group (Figure 3). Specifically, for each convolutional layer, we extracted channel-wise activation maps for all images in each dataset group. Let *a_ℓci_* denote the activation pattern for layer *ℓ*, channel *c*, and image *i*, vectorized over spatial positions. For each dataset group *k*, the raw stability score *r_ℓck_* was computed as the average cosine similarity over all ordered image pairs within that group:

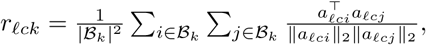

**Figure 3:**
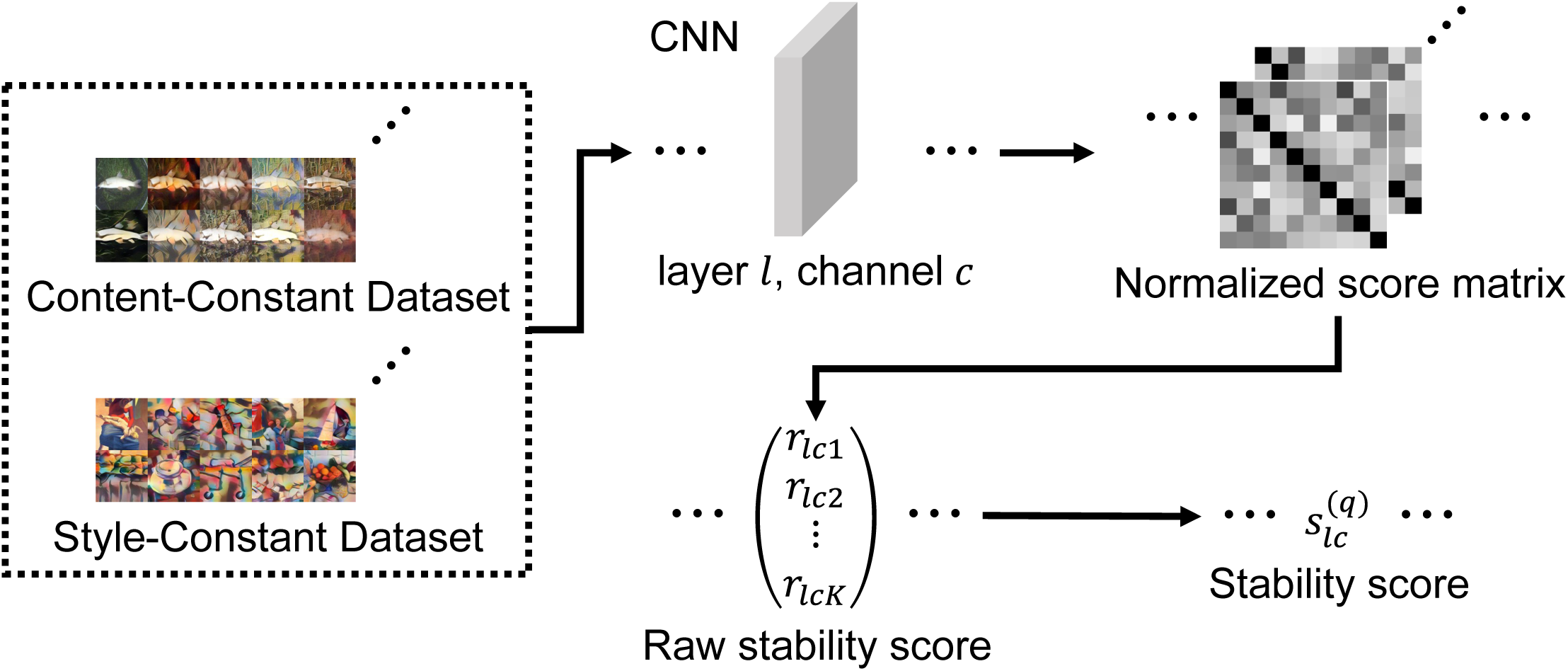
Computation of the Content Stability Score and Style Stability Score. For each convolutional layer and channel, activation maps were extracted for all images in each stimulus class and vectorized over spatial positions. Pairwise cosine similarities were then computed between the channel-wise activation patterns of all ordered image pairs within the stimulus group. Because each group comprised 10 images, this procedure produced a 10 *×* 10 similarity matrix. Averaging all entries of this matrix yielded a group-level raw stability score. The raw scores obtained from the 1,000 stimulus groups were subsequently combined using an aggregation criterion—mean, median, maximum, minimum, or entropy—to obtain one final stability score for each channel. Applying this procedure to the content-constant dataset yielded the Content Stability Score (CSS), whereas applying the same procedure to the style-constant dataset yielded the Style Stability Score (SSS). All images shown in this figure were generated by applying style transfer to images from ImageNet and WikiArt.

where *B_k_*denotes the set of images in group *k*.

These raw scores *r_ℓck_* were then aggregated across dataset groups to obtain a final channel score. Aggregation was necessary because each group provided a local estimate of response stability for a particular fixed content image or fixed style image, whereas pruning required one score per channel:

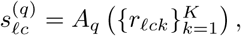

where *K* is the number of dataset groups and *A_q_* denotes the aggregation criterion. We considered several aggregation criteria, including average, median, maximum, minimum, and entropy across dataset groups. Using this two-step procedure, we computed two response-stability scores: the Content Stability Score and the Style Stability Score. The two scores were computed using the same raw-score and aggregation procedures, and differed only in the dataset from which the group-level scores were obtained. The Content Stability Score was computed from the content-constant dataset, whereas the Style Stability Score was computed from the style-constant dataset.

#### Score-based structured pruning

Using the response-stability scores described above, we performed structured pruning of convolutional output channels (Li, Kadav, Durdanovic, Samet, & Graf, 2016). In this pruning procedure, channels were ranked according to a channel-level importance measure, and a fixed proportion of low-importance channels was pruned according to a predefined pruning rate. This allowed us to construct submodels that preferentially retained channels consistent with each of the two candidate strategies.

Rather than ranking channels solely by their response-stability scores, we defined a score-modulated importance measure that also incorporated the L1 norm of the convolutional weights associated with each output channel. This hybrid criterion was intended to retain channels that were both stable under the target stimulus manipulation and non-negligible in the original network weights. Because pruning was performed within each convolutional layer, response-stability scores were first normalized separately for each layer. Let 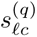 denote the original response-stability score for layer *ℓ*, channel *c*, and aggregation rule *q*. The normalized score 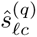 was defined as

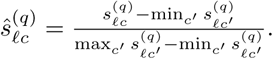

We then combined the normalized response-stability score with the L1 norm of the convolutional weights for each output channel. Let *W_ℓc_*denote the convolutional weights associated with output channel *c* in layer *ℓ*. The score-modulated channel importance was defined as

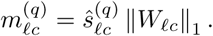

For each convolutional layer, channels were ranked according to 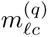. Given a pruning rate *p*, the lowest-scoring proportion *p* of output channels in that layer was pruned by setting all convolutional weights associated with those channels to zero. The first convolutional layer in the forward pass was left unpruned, and the specified pruning rate was applied to all remaining convolutional layers. Pruning was performed separately for each score type (Content Stability or Style Stability), aggregation rule, model architecture, and pruning rate. Because pruning alone substantially degraded classification performance, we fine-tuned each pruned submodel to retain viable object-recognition performance while preserving its channel-selection constraint.

#### Evaluation of submodels

We evaluated the submodels for two purposes. First, we aimed to characterize how the two pruning strategies, based on the Content Stability Score and the Style Stability Score, affected object-recognition performance. Second, because the raw stability scores were computed for individual stimulus groups and then aggregated across groups, we evaluated how different aggregation rules affected the performance of the resulting pruned models.

Models were evaluated using three sets of metrics. First, standard object-recognition performance was measured on the ImageNet validation set using top-1 classification accuracy. Second, robustness to stylized-image distribution shift was measured on the Stylized ImageNet test set, also using top-1 classification accuracy. Third, consistency with human behavioral responses was evaluated using model-vs-human benchmarks (Geirhos et al., 2021). For the model-vs-human benchmarks, we report model top-1 accuracy, accuracy difference, error consistency, and shape bias. Accuracy difference measured the gap between model and human top-1 accuracy. Error consistency measured whether models and humans tended to make errors on the same images. Shape bias was computed from texture– shape cue-conflict responses as the proportion of shape-consistent classifications among trials classified according to either the shape or texture label (Geirhos et al., 2019).

To summarize performance across pruning rates, we computed the normalized area under the pruning–performance curve (AUC). Models were evaluated at pruning rates from 0% to 100% in 10-percentage-point increments, and AUC was calculated using trapezoidal integration with pruning rates expressed as proportions between 0 and 1. For each pruning strategy and evaluation metric, AUC was first computed separately for each aggregation rule and then averaged across the five rules—mean, median, maximum, minimum, and entropy. This aggregation-averaged AUC was used to compare the overall performance of CSS- and SSS-based pruning.

To quantify the uncertainty in the aggregation-averaged AUC estimates, we performed a bootstrap analysis. For ImageNet and Stylized ImageNet, each bootstrap replicate consisted of 50,000 images sampled with replacement from the corresponding validation set. For the model-vs-human benchmarks, each replicate consisted of 18,080 stimulus identifiers, defined by dataset, condition, and filename, sampled with replacement from the full set of 18,080 stimuli. The model-vs-human metrics were recomputed from each resample; for shape bias, only eligible cue-conflict trials within the resample contributed to the calculation. Within each evaluation set, the same resample was shared across CSS- and SSS-based pruning, all five aggregation rules, and all pruning rates. The four model-vs-human metrics also shared the same resample. We obtained aggregation-averaged normalized AUCs for each pruning strategy over 1,000 bootstrap replicates.

Here, we focus on fine-tuned VGG-19, the architecture used for the subsequent image-synthesis experiment. The raw pruning-rate– metric-performance curves for the combinations of other architectures and fine-tuning stages are reported in Appendix: Pruned-model validation across architectures before and after fine-tuning.

Figure 4 shows the aggregation-averaged AUCs for the fine-tuned VGG-19 submodels. The CSS- and SSS-based pruning strategies differed significantly across all six evaluation metrics in two-sided null-centered paired-bootstrap tests after Bonferroni correction for the six metric-wise comparisons (all corrected *p < .*05). However, the absolute differences in normalized AUC were modest, with a maximum difference of only 0.026, despite their statistical significance. For a clearer visualization of these differences, the same results are presented with zoomed-in y-axis ranges in Appendix Zoomed-In Comparison of Normalized AUCs.

**Figure 4:**
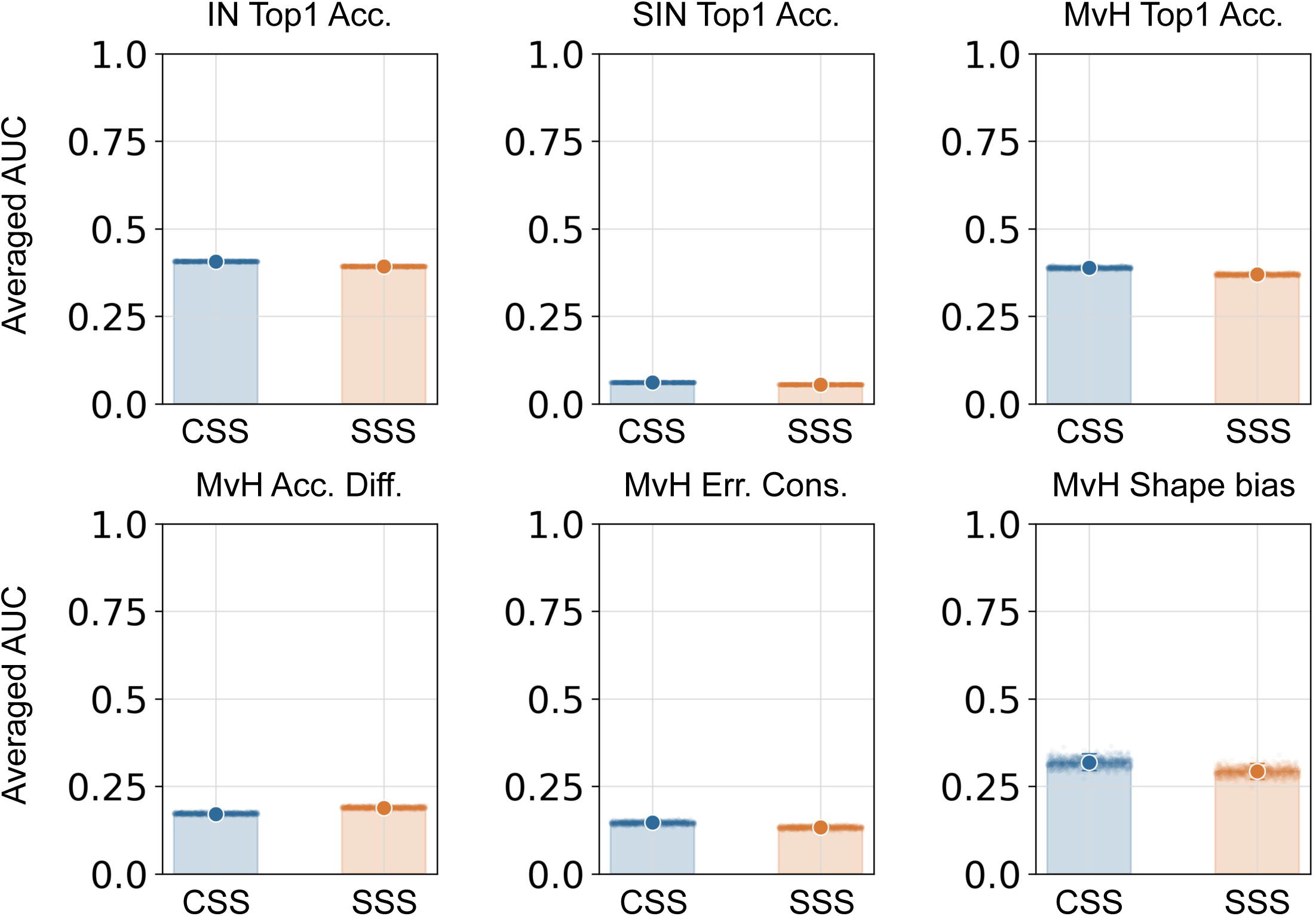
Comparison of CSS- and SSS-based pruning for fine-tuned VGG-19. From left to right, the top row shows ImageNet top-1 accuracy (IN Top1 Acc.), Stylized ImageNet top-1 accuracy (SIN Top1 Acc.), and model-vs-human top-1 accuracy (MvH Top1 Acc.); the bottom row shows accuracy difference (MvH Acc. Diff.), error consistency (MvH Err. Cons.), and shape bias (MvH Shape bias). Within each panel, the CSS-based submodel is shown on the left and SSS-based submodel on the right. Large points indicate full-sample estimates of normalized AUC across pruning rates from 0% to 100%, averaged across the five aggregation rules. Error bars indicate pointwise percentile 95% confidence intervals from 1,000 shared-resample bootstrap replicates, and translucent points show the 1,000 bootstrap estimates, using the same replicate indices for CSS and SSS. Due to the 0–1 y-axis range, the confidence intervals are too narrow to be visually discernible for the five metrics other than shape bias. CSS- and SSS-based submodels differed significantly for all six evaluation metrics after Bonferroni correction for the six metric-wise comparisons (all corrected *p* < .05), although the absolute differences in AUC were small. Higher values indicate better performance except for accuracy difference, for which lower values indicate closer agreement with human accuracy.

We next examined whether the choice of aggregation rule affected performance within each pruning strategy (Table 1). For the Content Stability Score-based submodels, mean aggregation produced the best performance for four of the six evaluation metrics. In contrast, the Style Stability Score-based submodels showed a different pattern: minimum aggregation produced the best or tied-best performance across the evaluation metrics. After Bonferroni correction for the 20 pairwise comparisons within each metric, mean aggregation for CSS and minimum aggregation for SSS each yielded significantly more favorable AUCs than all four alternative rules for three of the six evaluation metrics (all corrected *p < .*05).

**Table 1:** Normalized area under the pruning-performance curve for the fine-tuned VGG-19. Values summarize each evaluation metric across pruning rates from 0% to 100% in 10-percentage-point increments. Results are shown separately for Content Stability Score (CSS)- and Style Stability Score (SSS)-based submodels using the mean (Avg.), median (Med.), maximum (Max), minimum (Min), and entropy (Ent.) aggregation rules. The evaluation metrics comprise top-1 classification accuracy on the ImageNet validation set (IN Top1 Acc.), top-1 classification accuracy on the Stylized ImageNet test set (SIN Top1 Acc.), model top-1 classification accuracy on the model-vs-human benchmark (MvH Top1 Acc.), accuracy difference between the model and humans (MvH Acc. Diff.), error consistency (MvH Err. Cons.), and shape bias (MvH Shape bias). Boldface indicates the best-performing aggregation rule within each submodel for each metric; the lowest value is considered best for accuracy difference, whereas the highest value is considered best for all other metrics.

| Metric | CSS Avg. | CSS Med. | CSS Max | CSS Min | CSS Ent. | SSS Avg. | SSS Med. | SSS Max | SSS Min | SSS Ent. |
| --- | --- | --- | --- | --- | --- | --- | --- | --- | --- | --- |
| IN Top1 Acc. | 0.420 | **0.422** | 0.391 | 0.399 | 0.402 | 0.386 | 0.392 | 0.372 | **0.411** | 0.402 |
| SIN Top1 Acc. | **0.066** | 0.065 | 0.062 | 0.059 | 0.052 | 0.056 | **0.057** | 0.051 | **0.057** | 0.054 |
| MvH Top1 Acc. | **0.408** | 0.406 | 0.361 | 0.390 | 0.376 | 0.371 | 0.372 | 0.350 | **0.392** | 0.360 |
| MvH Acc. Diff. | **0.152** | 0.155 | 0.202 | 0.166 | 0.182 | 0.182 | 0.182 | 0.205 | **0.166** | 0.206 |
| MvH Err. Cons. | **0.157** | 0.155 | 0.142 | 0.143 | 0.135 | 0.132 | 0.130 | 0.121 | **0.144** | 0.139 |
| MvH Shape bias | 0.344 | 0.334 | **0.348** | 0.303 | 0.261 | **0.301** | 0.291 | 0.277 | **0.301** | 0.292 |

## Discussion

The aggregation analysis revealed a systematic difference between the two pruning strategies. For the CSS-based submodels, mean aggregation yielded the best performance for four of the six evaluation metrics. Mean aggregation favors channels that exhibit high response stability on average across content-defined stimulus groups. By contrast, minimum aggregation yielded the best or tied-best performance across the evaluation metrics for the SSS-based submodels. Minimum aggregation provides a more conservative criterion by assigning each channel the score of its least stable style-defined stimulus group, thereby favoring channels whose responses remain stable across a broad range of styles. These findings suggest that the most appropriate method for summarizing group-level response stability depends on whether stability is evaluated across changes in visual style or changes in object content. Accordingly, mean aggregation for CSS and minimum aggregation for SSS were used to define the representative submodels for the subsequent stimulus-generation analysis.

Importantly, comparable recognition performance does not imply that the CSS- and SSS-based submodels encode visual information in the same way. The two pruning strategies were derived from complementary stimulus manipulations and preferentially retained different channel subsets. Their internal feature subspaces may therefore differ even when they support similar category-level outputs. The recognition and model-vs-human benchmarks used here may be insufficiently sensitive to reveal such representational differences. To test whether the two feature subspaces nevertheless impose different perceptual constraints, we next used the selected CSS- and SSS-based submodels to generate matched style-transferred images and conducted a behavioral experiment to determine which submodel-derived images more closely reflected the visual style of a reference image according to human observers.

## Experiment 2: Behavioral evaluation of synthesized stimuli

To examine the perceptual relevance of the two computational strategies, we next evaluated images synthesized using the corresponding submodel-derived feature subspaces. Model-metamer studies provide a useful general framework for this purpose. If a model representation captures dimensions that are important for human perception, images matched within that representation should preserve the corresponding perceptual attributes for human observers (Feather et al., 2023). Here, we applied this logic specifically to the perception of visual style.

Neural style transfer modifies a content image so that its model-defined style statistics approximate those of a target style image while preserving the structure of the content image. Accordingly, the synthesized images provide a visualization of the style constraints imposed by the selected model representation (e.g., Gram matrix). We generated matched style-transfer images from the CSS-based and SSS-based synthesis subspaces. We then conducted a behavioral experiment in which observers judged which generated image better reflected the target style. This comparison tested which of the two model-derived subspaces imposed constraints that were more consistent with human judgments of visual style.

## Methods

### Stimulus generation

For style transfer, we used the Gram-matrix-based neural style transfer method proposed by Gatys, Ecker, and Bethge (2016). Although this method can be applied to a wide range of convolutional neural networks, we used VGG-19 submodels to remain consistent with the original formulation and with the submodel construction described in the previous section. For the final behavioral stimulus set, we used the 30% pruning condition, because this pruning level preserved usable object-recognition performance while still yielding distinct submodel-derived feature subspaces.

Among the CSS-based and SSS-based submodels constructed in the previous section, we further restricted the final synthesis comparison to channels that were exclusive to each submodel. Channels retained by both submodels were treated as shared channels, whereas channels retained by only one submodel were treated as exclusive channels. This procedure was used so that the behavioral comparison would reflect differences between the two computational strategies rather than properties of channels shared by both submodels.

For a generation condition *g*, let 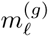 denote the binary channel mask applied to layer *ℓ*, and let 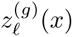 (*x*) denote the activation tensor of the corresponding pruned VGG-19 checkpoint for image *x*. The masked activation was defined as

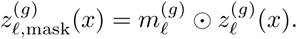

where *⊙* denotes element-wise multiplication. The first convolutional layer was treated as an exception: no channel mask was applied to this layer, equivalently setting 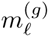 to an all-one mask.

In the final stimulus-generation procedure, the content loss was defined as the mean squared error between the content-layer features of the generated image and those of the content reference image. These features were computed using a fixed ImageNet-pretrained VGG-19 network without channel masking. The style loss was defined as the mean squared error between the Gram matrices of the generated image and the style reference image, computed from the layer-specific masked activations of the corresponding fine-tuned, pruned VGG-19 checkpoint. We also included a total-variation regularizer that penalized local pixel differences (Mahendran & Vedaldi, 2015). Because the content loss was computed identically across generation conditions, whereas the model-derived feature subspace used for the style loss differed between conditions, the experimental manipulation was restricted to the style-feature subspace. The total loss was

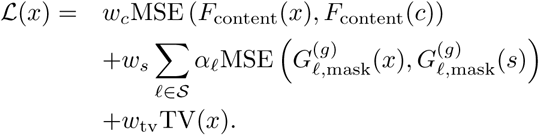

Here, *x* denotes the generated image, *c* the content reference image, and *s* the style reference image. *F*_content_(*·*) denotes the content-layer feature from the fixed ImageNet-pretrained VGG-19 network, and 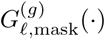 denotes the Gram matrix computed from 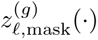. The Gram matrix was normalized by (*C_ℓ_H_ℓ_W_ℓ_*)*^−^*^1^. The hyperparameters were set to *w_c_* = 0.5, *w_s_* = 7.5 *×* 10^7^, and *w*_tv_ = 10*^−^*^7^; the style-layer weights *α_ℓ_* were 0.25 for conv1 layers, 0.5 for conv2 layers, 2.0 for conv3 layers, 4.0 for conv4 layers, and 8.0 for conv5 layers.

### Stimulus sampling, participants, and apparatus

Candidate stimuli were generated from all combinations of five content images selected from the ImageNet validation set based on semantic diversity across classes and an image-quality score incorporating contrast, brightness, aspect ratio, and the relative strength of edges near the image center and 500 style images randomly sampled from the WikiArt dataset. For each content–style combination, we generated a matched pair of synthesized images using the CSS-based and SSS-based synthesis subspaces. Candidate pairs were manually excluded if either synthesized image showed obvious synthesis artifacts.

From the remaining candidates, style images for the behavioral experiment were selected using LPIPS-Alex, a perceptual image-distance metric (Zhang, Isola, Efros, Shechtman, & Wang, 2018). To avoid selecting stimuli solely on the basis of one synthesis condition, we selected equal numbers of styles using rankings computed separately for the CSS-based and SSS-based conditions. For each condition, we computed the mean LPIPS distance between the generated images and their corresponding content references across the five content images. Styles were then ranked according to the perceptual dissimilarity of the generated images from the content references. We selected the five styles whose generated images were least similar to the content references in the CSS-based condition and five additional, non-overlapping styles from the corresponding SSS-based ranking. This balanced selection procedure allowed the stimulus set to include styles that produced pronounced image changes under each of the two synthesis conditions. Pairing each of these 10 styles with each of the five content images yielded 50 content–style conditions for the behavioral experiment (Figure 5). An additional control analysis examining the effect of the aggregation rule on image synthesis is reported in Appendix: Control analysis of aggregation-rule effects on image synthesis.

**Figure 5:**
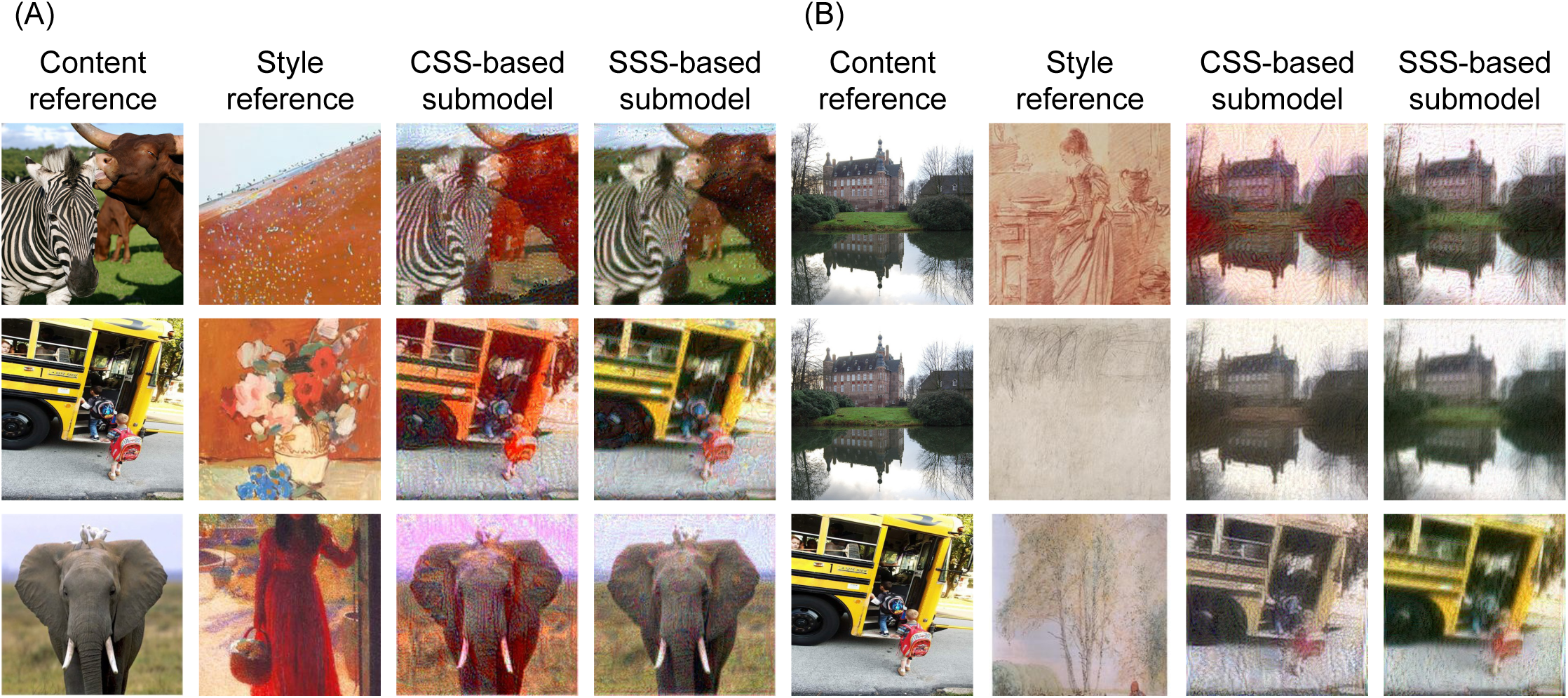
Representative examples from the final stimulus set. Within each panel, columns show the ImageNet content reference, WikiArt style reference, and the corresponding images generated using the CSS- and SSS-based synthesis subspaces, respectively. (A) Content–style pairs selected according to the LPIPS-Alex distance between the CSS-generated image and its content reference. The pairs producing the five largest distances among the 500 candidates were selected. (B) Content–style pairs selected using the same procedure applied to the SSS-generated images. All images shown in this figure were either obtained from ImageNet or WikiArt, or generated by applying style transfer to images from these sources.

Ten observers completed the behavioral experiment. All observers had normal or corrected-to-normal vision and provided informed consent before participation. The study was conducted with approval by the Ethics and Safety Committee of the University of Tokyo.

Stimuli were presented on a 24.5-inch LCD calibrated display (XL2546X) against a dark gray background. Each image was displayed within a 5-cm square frame at a viewing distance of 57 cm, subtending approximately 5*^◦^ ×* 5*^◦^* of visual angle.

### Procedure

We used a two-alternative forced-choice (2AFC) judgment task of style similarity. On each trial, observers viewed an ImageNet content reference (Figure 6 (A) top), a WikiArt style reference (Figure 6 (A) bottom), and two synthesized images generated from the same content–style pair (Figure 6 (A) left and right). Observers were instructed to indicate by button press which synthesized image had a visual style more similar to that of the style reference image. The two alternatives were generated using the CSS-based and SSS-based synthesis subspaces, respectively. Their left–right positions were randomized on each trial.

**Figure 6:**
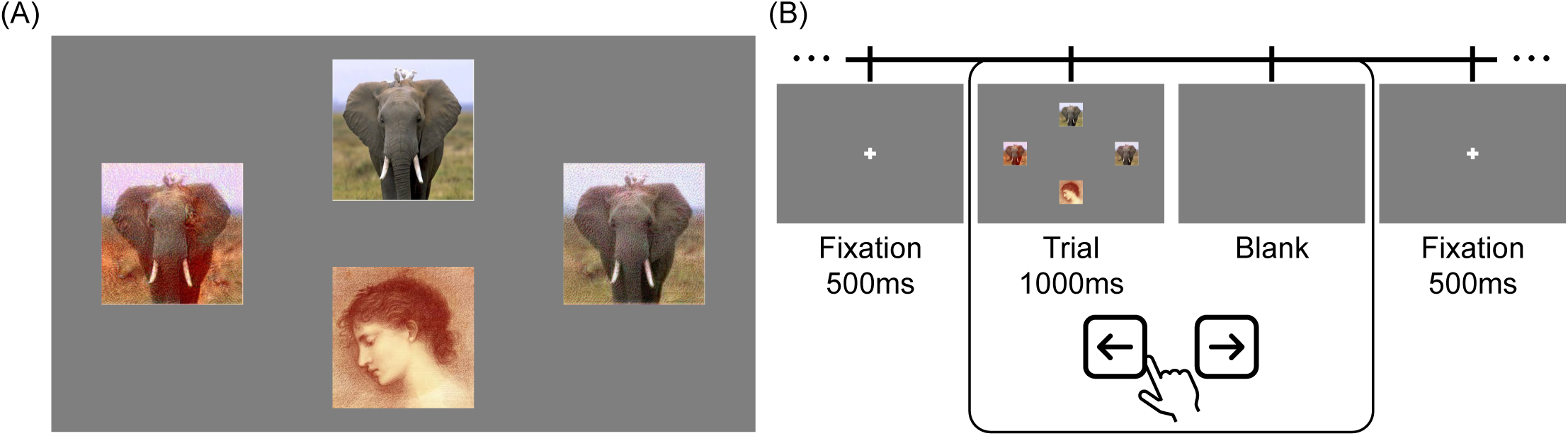
Overview of the experimental procedure. (A) Stimulus display. The content image was presented at the top and the style reference image at the bottom. The two images presented on the left and right were stylized images generated using the CSS- and SSS-based subspaces, with their positions randomized across trials. (B) Trial sequence. A fixation screen was presented for 500 ms, followed by the stimulus display for 1,000 ms. A blank screen was then presented until the observer responded. Observers indicated by button press which of the two synthesized images was more similar in visual style to the style reference image. This sequence was repeated for each trial. All images shown in this figure were either obtained from ImageNet or WikiArt, or generated by applying style transfer to images from these sources.

Each trial began with a fixation cross presented for 500 ms, followed by the four-image stimulus display for 1000 ms. The stimulus display was then replaced by a blank gray screen, which remained visible until the observer responded (Figure 6 (B)). Responses and reaction times were recorded on each trial.

Each of the 50 content–style conditions was presented 10 times, resulting in 500 trials per observer. The resulting trials were presented in random order, and observers were given a break after every 100 trials.

Trials with reaction times shorter than 200 ms or longer than 5000 ms were designated for exclusion, although no trials met these criteria. Additional reaction-time analyses are reported in Appendix: Behavioral follow-up analyses.

Written informed consent was obtained from all participants prior to participation. The study protocol was approved by the institutional ethics committee of the Graduate School of Information Science and Technology, The University of Tokyo (approval date: February 10, 2026; reference number: E2025ALS220-01). All procedures were conducted in accordance with the Declaration of Helsinki and relevant institutional guidelines and regulations.

## Results and Discussion

Trial-level choices were analyzed using a generalized linear mixed-effects model with a Bernoulli likelihood and a logit link. Selection of the CSS-based image was coded as 1 and selection of the SSS-based image as 0. For participant *i*, content *j*, style *k*, and repeated trial *l*, the model was specified as

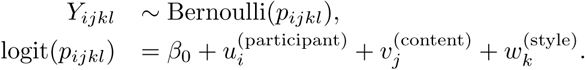

where *β*_0_ denotes the fixed intercept and *u_i_*, *v_j_*, and *w_k_* denote random intercepts for participant, content, and style, respectively. Because *β*_0_ = 0 corresponds to a CSS-choice probability of 0.5, the fixed intercept was tested against zero using a two-sided Wald test.

Across 5,000 valid trials, the observed CSS-based response rate was 0.746. The mixed-effects logistic regression revealed a significantly positive fixed intercept, *β*_0_ = 1.30, *SE* = 0.28, *z* = 4.65, *p < .*001, indicating that CSS-based images were selected more often than chance after accounting for variability across participants, contents, and styles. The corresponding model-estimated CSS-choice probability was 0.786 (95% CI [0.679, 0.864]).

To complement this trial-level analysis, we also summarized CSS-based response rates at the participant level and compared them with chance. For each observer, we calculated the CSS-based response rate, defined as the proportion of trials in which the CSS-based image was judged to be more similar in visual style to the reference image. Participant-level response rates were compared with chance level (0.5) using a two-sided one-sample *t*-test, with a two-sided Wilcoxon signed-rank test as a non-parametric robustness check.

The mean CSS-based response rate across all conditions was 0.746 (95% confidence interval: 0.657–0.835; Figure 7 (A)), which was significantly above chance, *t*(9) = 6.23, *p < .*001. The Wilcoxon signed-rank test yielded the same conclusion, *p* = .004, indicating that observers more often judged images generated using the CSS-based subspace as more similar in visual style to the reference image.

**Figure 7:**
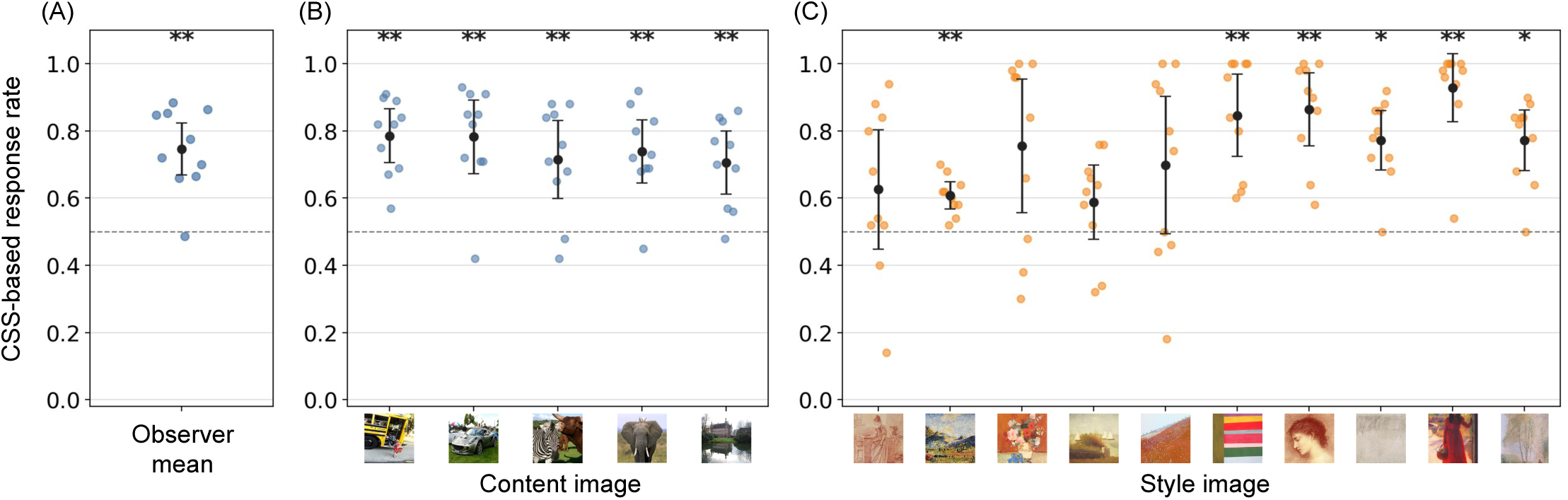
CSS-based response rates in the 2AFC style-similarity task. (A) Mean response rate across all conditions. (B) Content-wise response rates. (C) Style-wise response rates. Points indicate participant-level response rates, and error bars indicate 95% confidence intervals across participants. The dashed line denotes chance level (0.5). All images shown in this figure were either obtained from ImageNet or WikiArt.

This preference was observed for each of the five content images (two-sided Wilcoxon signed-rank tests, Bonferroni-corrected separately across content images: all *p*_Bonferroni_ *< .*05; Figure 7 (B)). At the level of individual style images, however, the magnitude of the preference varied across styles (Figure 7 (C)). Specifically, images generated using the CSS-based subspace were selected significantly more often for 6 of the 10 styles (two-sided Wilcoxon signed-rank tests, Bonferroni-corrected separately across style images: all *p*_Bonferroni_ *< .*05), whereas response rates for the remaining 4 styles did not differ significantly from chance (all *p*_Bonferroni_ *> .*1). Importantly, no style showed a significantly below-chance CSS-based response rate, indicating no significant preference for images generated using the SSS-based subspace.

Taken together, these results suggest that the CSS-based synthesis subspace produced images that were more consistent with human judgments of visual-style similarity than those produced by the SSS-based subspace.

## General Discussion

The present study asked whether computational subspaces that support similar style-robust object-recognition performance are equally consistent with human judgments of visual style. We compared two candidate strategies within neural networks trained on Stylized ImageNet. CSS-based submodels retained channels whose responses were stable across style changes when content was fixed. SSS-based submodels retained channels whose responses were stable across content changes when style was fixed. The two submodel families performed similarly on ImageNet, Stylized ImageNet, and model–vs-human benchmarks. We then used feature channels exclusive to each VGG-19 submodel to constrain Gram-matrix-based style transfer. Observers judged images generated from the CSS-based subspace as more similar in visual style to the reference. These results show that the CSS-based subspace was more consistent with human style judgments than the SSS-based subspace.

An implication of the present findings is that similarity in task-level performance does not guarantee similarity in perceptually relevant internal computation. The differences between the CSS- and SSS-based submodels were modest on object recognition and model-vs-human benchmarks, yet their synthesized images produced markedly different human judgments. Related dissociations have also been reported for recent foundation models. For example, CLIP and related vision-language models can reproduce selected human-like patterns of semantic or language-biased judgment, while detailed analyses of their representations have revealed substantial departures from the organization of human cognitive processes (e.g., Lemesle et al., 2022; Tschannen, Mustafa, & Houlsby, 2023). Resemblance at the level of behavioral decisions or neural-response predictions should therefore be regarded as a starting point for evaluating human-model alignment, rather than as evidence that the underlying mechanisms are shared.

Our finding that images generated using the CSS-based subspace were more consistent with human style judgments may be counterintuitive at first glance. The CSS-based submodel retained channels whose spatial activation patterns remained relatively stable when the same content was rendered in different styles. Therefore, one might expect these channels to attenuate or discard style variation. Nevertheless, images constrained by the exclusive CSS-based channels conveyed the reference style more effectively than those constrained by the SSS-based channels. One possible explanation lies in the difference between the criterion used to select channels and the representation used for image synthesis. CSS was computed separately for each channel from the similarity of its activation maps, whereas the style-transfer objective was based on Gram matrices, which capture second-order relationships among multiple channel responses. High stability at the level of individual channels therefore does not necessarily imply an absence of style information in their multivariate correlation structure. A collection of content-stable channels may provide a stable representational basis, while style is expressed through relative relationships among those channels.

This potential division between a stable representational basis and complementary variation is conceptually similar to findings in shape and material perception. Variations in object materials can act as perturbations that interfere with shape estimation. Nevertheless, images contain shape cues that are robust to material variation (e.g., image-gradient direction/orientation), and humans are known to use these cues to estimate object shape (Srinath et al., 2021). Conversely, after image-gradient direction/orientation has been used as a material-invariant cue for shape estimation, variation in image-gradient magnitude remains as complementary information that supports human material perception (Sawayama & Nishida, 2018; Nishida & Sawayama, 2026). From the perspective of object recognition, variations in style and material can both be regarded as noise-like variations that disrupt stable recognition. The present findings raise the possibility that acquiring representations robust to such variations may, paradoxically, enable the visual system to richly encode the residual information associated with those same variations.

The present approach is also related to the use of model perturbations and synthesized stimuli to compare artificial and biological vision. In the model-metamer literature, images are optimized to match the internal representation of a reference image and are subsequently tested for perceptual equivalence in human observers. Here, our approach aims to improve model interpretability by constructing submodels that instantiate distinct hypothesized computational strategies and testing which submodel generates stimuli more consistent with human perception. This strategy also differs from approaches that begin by constructing compact models, although neither approach can be considered generally superior. Compactness alone does not necessarily ensure interpretability, but compact-model approaches have nevertheless achieved notable successes (e.g., Cowley, Stan, Pillow, & Smith, 2026; Morimoto et al., 2026). A strength of our approach is that submodels can be constructed to instantiate explicit a priori hypotheses. Conversely, its conclusions are necessarily constrained by those hypotheses and may fail to identify the relevant computations if the hypotheses are poorly specified. A mechanistic understanding of deep models will therefore likely require converging evidence from multiple complementary approaches.

The present study also has several limitations. The behavioral experiment provided a relative comparison between two candidate synthesis subspaces rather than an absolute measure of human–model alignment. The above-chance preference for CSS-based images indicates that, under the present procedure, this subspace produced style-transfer constraints that were more consistent with human judgments than those produced by the SSS-based subspace. This finding neither establishes that the CSS-based subspace fully captures human representations of visual style nor demonstrates that the SSS-based subspace contains no perceptually relevant style information. Further research is therefore needed to quantify the extent to which each subspace captures human style perception.

In summary, the present study provides a framework for moving from performance-level alignment toward experimentally testable hypotheses about internal computation. By isolating candidate subspaces within a common parent model, expressing their computational consequences through image synthesis, and evaluating those images with human observers, the approach can reveal perceptually meaningful differences that remain hidden in standard benchmarks. In the case of style-invariant object recognition, the results identify representations that remain stable across changes in visual style as a particularly promising basis for understanding how object content and visual style may be jointly represented.

## Code and Data Availability

Code and de-identified behavioral response data are being prepared for public release and are not yet publicly available.

## Acknowledgments and Disclosure

This work was supported by JST Moonshot R&D Program, Grant Number JPMJMS259G, JST SPRING, Grant Number JP-MJSP2108, and JSPS KAKENHI Grant Numbers 26K02927 and 26K03250.

OpenAI’s ChatGPT was used for editing and coding assistance. All scientific content, analyses, and conclusions were verified by the authors.

## Appendix

### Fine-tuning on Stylized ImageNet

We evaluated fine-tuning on Stylized ImageNet as a validation step for the models subsequently used for score-based channel pruning. This appendix analysis supports the model-construction pipeline described in Construction of computational submodels by confirming, prior to the computation of response-stability scores, that the models responded meaningfully to stylized images rather than only to the original ImageNet distribution. The aim was not to provide a new evaluation of Stylized ImageNet training, but to verify the models used for subspace extraction retained adequate object recognition performance after being fine-tuned on stylized images.

For all tested architectures, validation accuracy on Stylized ImageNet increased rapidly during the first few epochs and subsequently reached a broad plateau (Figure 8 (A)). For each architecture, the checkpoint with the highest validation accuracy across the 50 training epochs was selected for the computation of response-stability scores. ResNet-50 achieved the highest validation accuracy at its selected checkpoint, followed by CORnet-S, VGG-19, and AlexNet. Specifically, the selected checkpoints achieved top-1 accuracies of 0.3357 for AlexNet at epoch 23, 0.5084 for CORnet-S at epoch 7, 0.5632 for ResNet-50 at epoch 7, and 0.4467 for VGG-19 at epoch 12. The VGG-19 checkpoint selected at epoch 12 was subsequently used in the final stimulus-generation pipeline.

**Figure 8:**
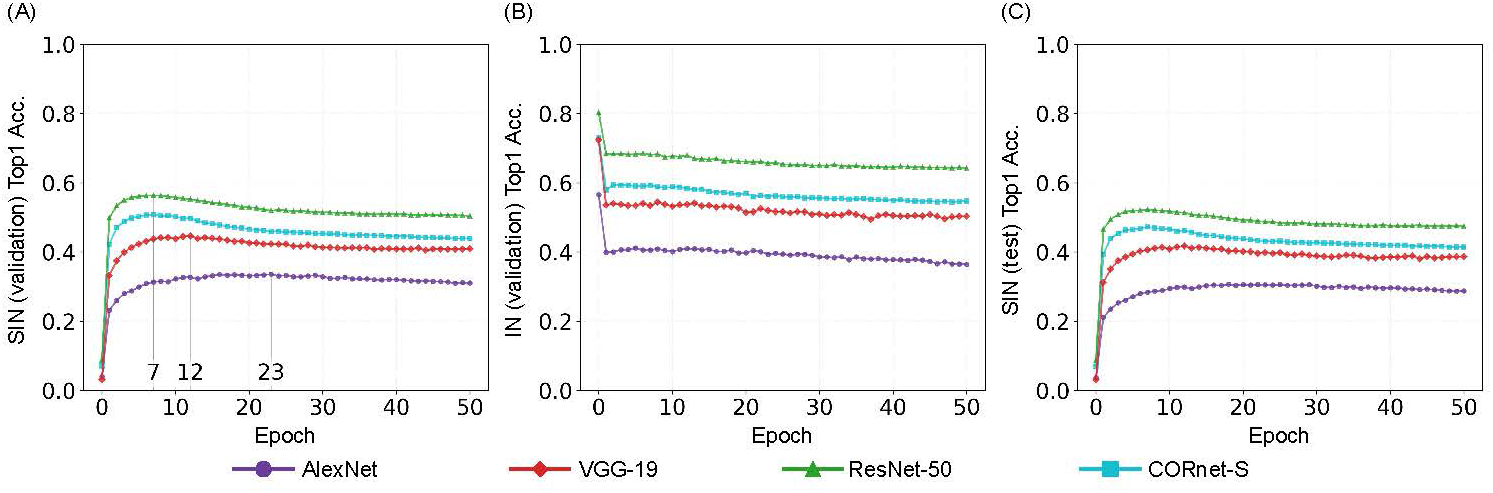
Object-recognition performance of models fine-tuned on Stylized ImageNet. (A) Top-1 accuracy on the Stylized ImageNet validation set, generated by stylizing a subset of the ImageNet training set. (B) Top-1 accuracy on the original ImageNet validation set. (C) Top-1 accuracy on the Stylized ImageNet test set, generated by stylizing the ImageNet validation set. Each panel shows the performance of all four candidate architectures: AlexNet, VGG-19, ResNet-50, and CORnet-S.

Accuracy on the original ImageNet validation set decreased during the first few epochs of fine-tuning (Figure 8 (B)). By contrast, accuracy on the Stylized ImageNet test set, generated by stylizing images from the ImageNet validation set, increased in a manner similar to that observed on the Stylized ImageNet validation set (Figure 8 (C)).

### Pruned-model validation across architectures before and after fine-tuning

We evaluated the performance of the pruned models across architectures, pruning rates, aggregation criteria, score-calculation methods, and fine-tuning stages. This analysis was intended to characterize the broader behavior of the score-based pruning and to verify that the validation results obtained with the fine-tuned VGG-19 were not specific to a single architecture or experimental setting. Here, we report systematic pruning sweeps for AlexNet, VGG-19, ResNet-50, and CORnet-S, complementing the fine-tuned VGG-19 analysis reported in Evaluation of submodels. These supplementary analyses provide broader context for the pruning regime from which the VGG-19 submodels used for stimulus generation were selected.

Across architectures, post-fine-tuning models generally preserved performance over a broader range of pruning rates than the corresponding pre-fine-tuning models. This pattern was particularly evident for AlexNet and ResNet-50, for which score-based pruning exhibited more gradual performance degradation after fine-tuning than before fine-tuning. In contrast, pre-fine-tuning models often showed substantial degradation even at relatively low pruning rates, suggesting that fine-tuning mitigated part of the functional loss associated with channel removal. Content Stability Score (CSS)- and Style Stability Score (SSS)-based pruning also showed broadly comparable degradation profiles across many evaluation metrics, although their relative performance varied with architecture, pruning rate, aggregation criterion, score-calculation method, evaluation metric, and fine-tuning stage (Figures 9–12). Thus, the pruning sweeps are better interpreted as a model-selection and validation procedure than as evidence for a single, consistently superior pruning criterion. The fine-tuned VGG-19 submodels used for stimulus generation were selected from a pruning regime in which measurable object-recognition and model-vs-human benchmark performance was retained, while the broader sweeps reported here establish the architectural and methodological context for that selection.

**Figure 9:**
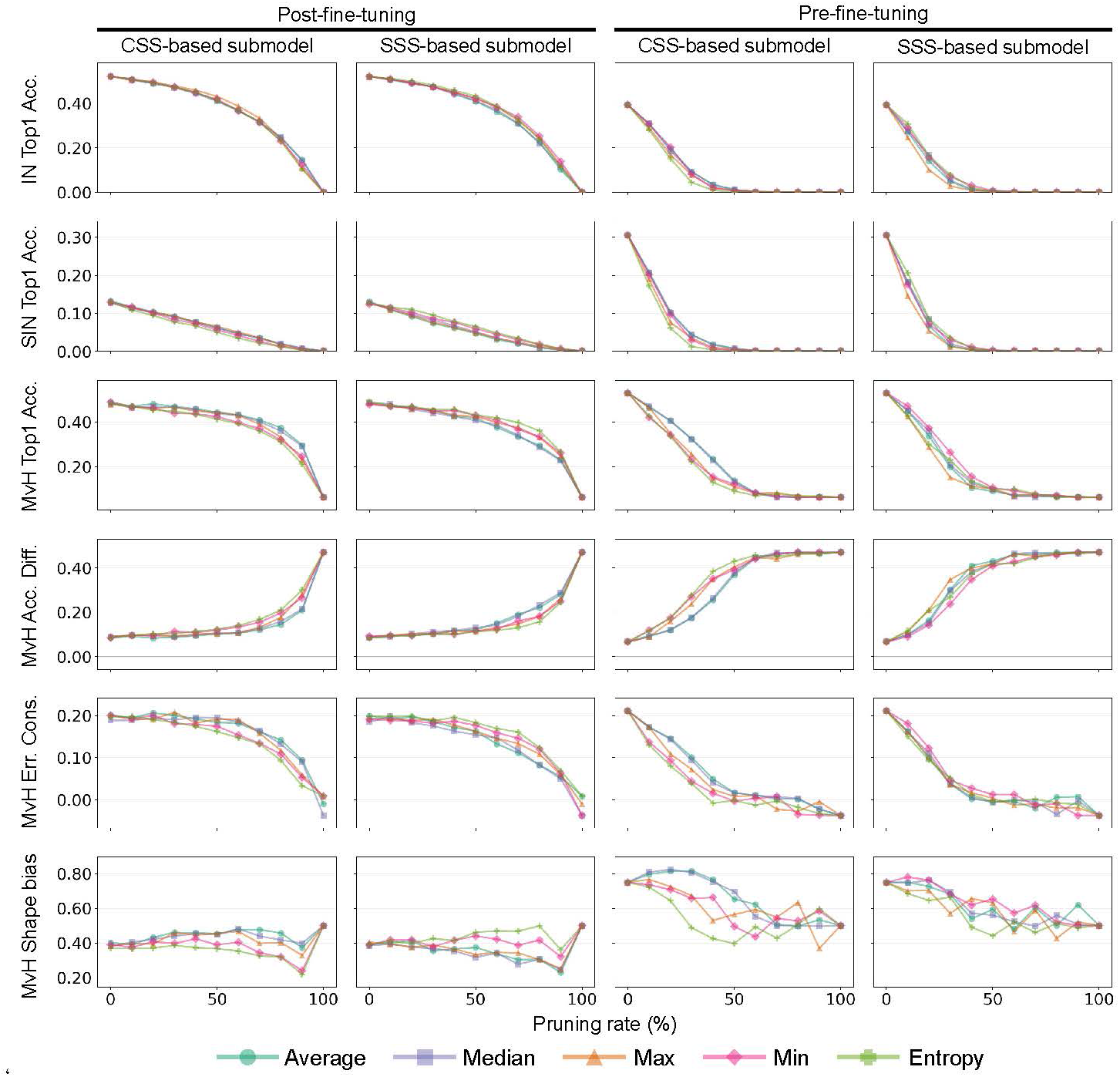
AlexNet performance under score-based pruning. Rows show evaluation metrics. From top to bottom, these are Top-1 classification accuracy on ImageNet (IN Top1 Acc.), Top-1 classification accuracy on Stylized ImageNet (SIN Top1 Acc.), and four Model-vs-Human (MvH) metrics: Top-1 classification accuracy (MvH Top1 Acc.), the accuracy difference between the model and human observers (MvH Acc. Diff.), error consistency (MvH Err. Cons.), and shape bias (MvH Shape bias). Columns show combinations of score-calculation method and fine-tuning stage. Curves show performance as a function of pruning rate for different aggregation criteria.

**Figure 10:**
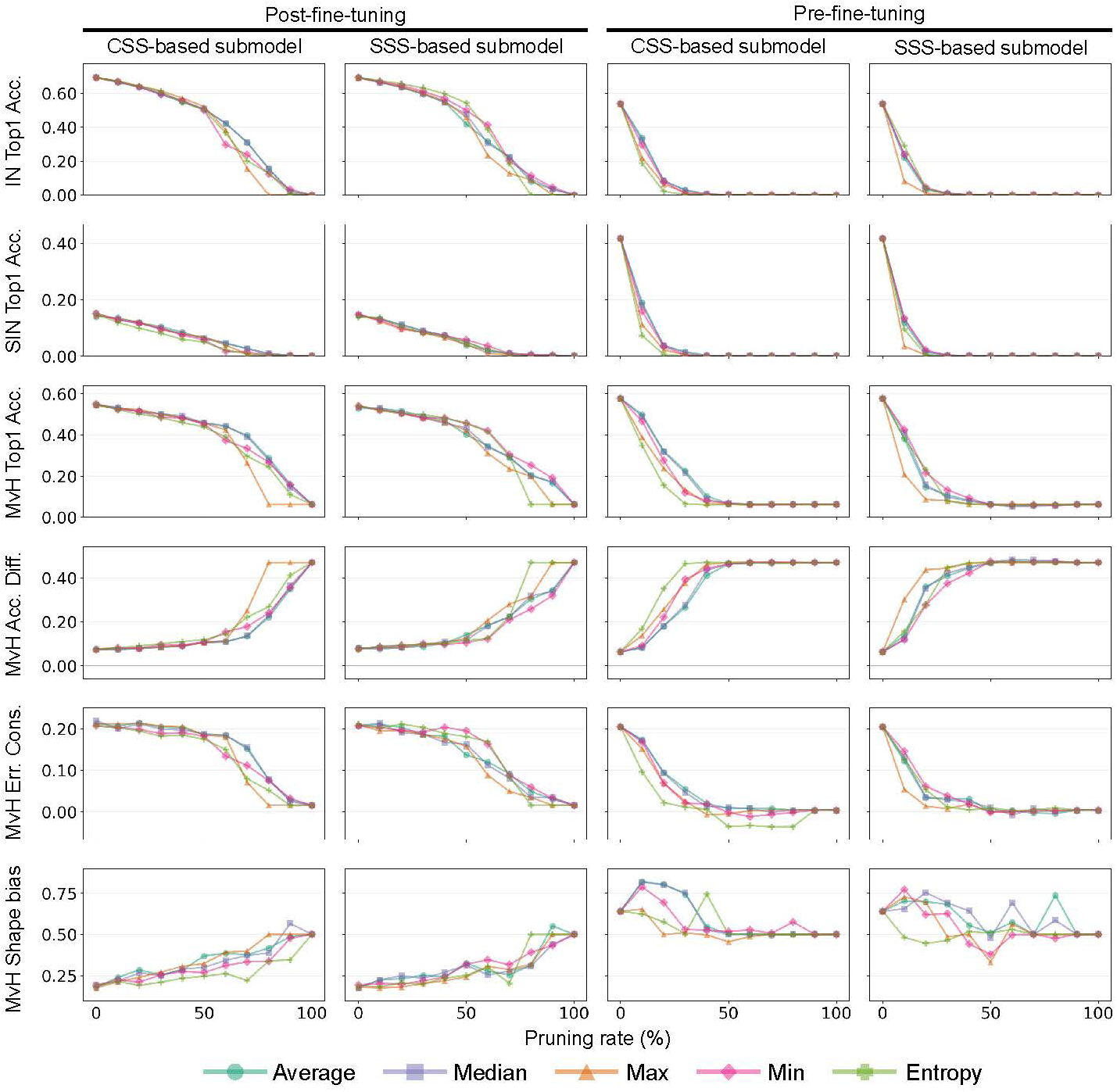
VGG-19 performance under score-based pruning. Rows show evaluation metrics. From top to bottom, these are Top-1 classification accuracy on ImageNet (IN Top1 Acc.), Top-1 classification accuracy on Stylized ImageNet (SIN Top1 Acc.), and four Model-vs-Human (MvH) metrics: Top-1 classification accuracy (MvH Top1 Acc.), the accuracy difference between the model and human observers (MvH Acc. Diff.), error consistency (MvH Err. Cons.), and shape bias (MvH Shape bias). Columns show combinations of score-calculation method and fine-tuning stage. Curves show performance as a function of pruning rate for different aggregation criteria.

**Figure 11:**
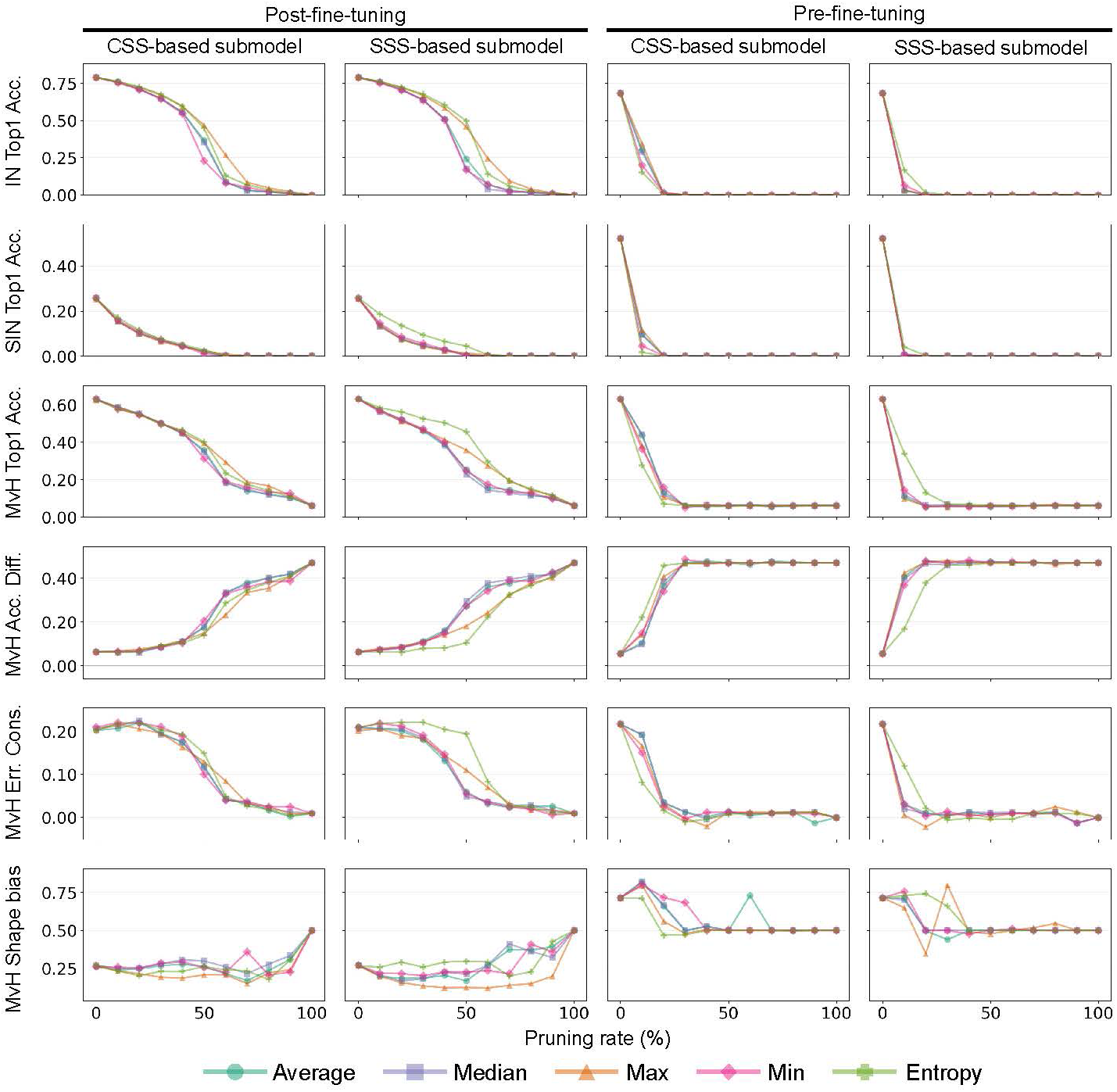
ResNet-50 performance under score-based pruning. Rows show evaluation metrics. From top to bottom, these are Top-1 classification accuracy on ImageNet (IN Top1 Acc.), Top-1 classification accuracy on Stylized ImageNet (SIN Top1 Acc.), and four Model-vs-Human (MvH) metrics: Top-1 classification accuracy (MvH Top1 Acc.), the accuracy difference between the model and human observers (MvH Acc. Diff.), error consistency (MvH Err. Cons.), and shape bias (MvH Shape bias). Columns show combinations of score-calculation method and fine-tuning stage. Curves show performance as a function of pruning rate for different aggregation criteria.

**Figure 12:**
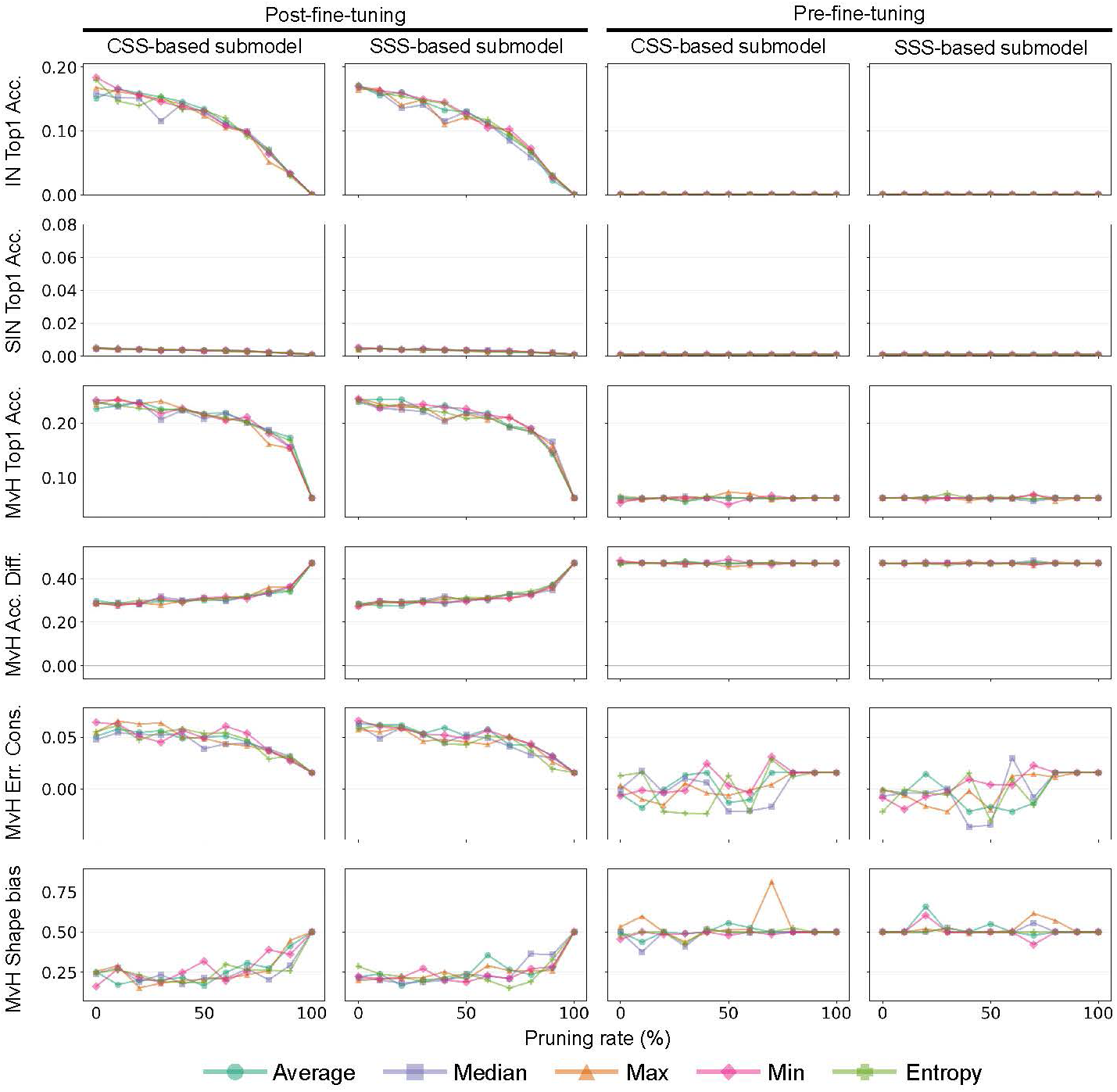
CORnet-S performance under score-based pruning. Rows show evaluation metrics. From top to bottom, these are Top-1 classification accuracy on ImageNet (IN Top1 Acc.), Top-1 classification accuracy on Stylized ImageNet (SIN Top1 Acc.), and four Model-vs-Human (MvH) metrics: Top-1 classification accuracy (MvH Top1 Acc.), the accuracy difference between the model and human observers (MvH Acc. Diff.), error consistency (MvH Err. Cons.), and shape bias (MvH Shape bias). Columns show combinations of score-calculation method and fine-tuning stage. Curves show performance as a function of pruning rate for different aggregation criteria.

### Zoomed-In Comparison of Normalized AUCs

Figure 13 presents the same aggregation-averaged normalized AUCs as Figure 4, but with zoomed-in y-axis ranges. This visualization makes the small differences between CSS- and SSS-based submodels and their bootstrap confidence intervals easier to discern.

**Figure 13:**
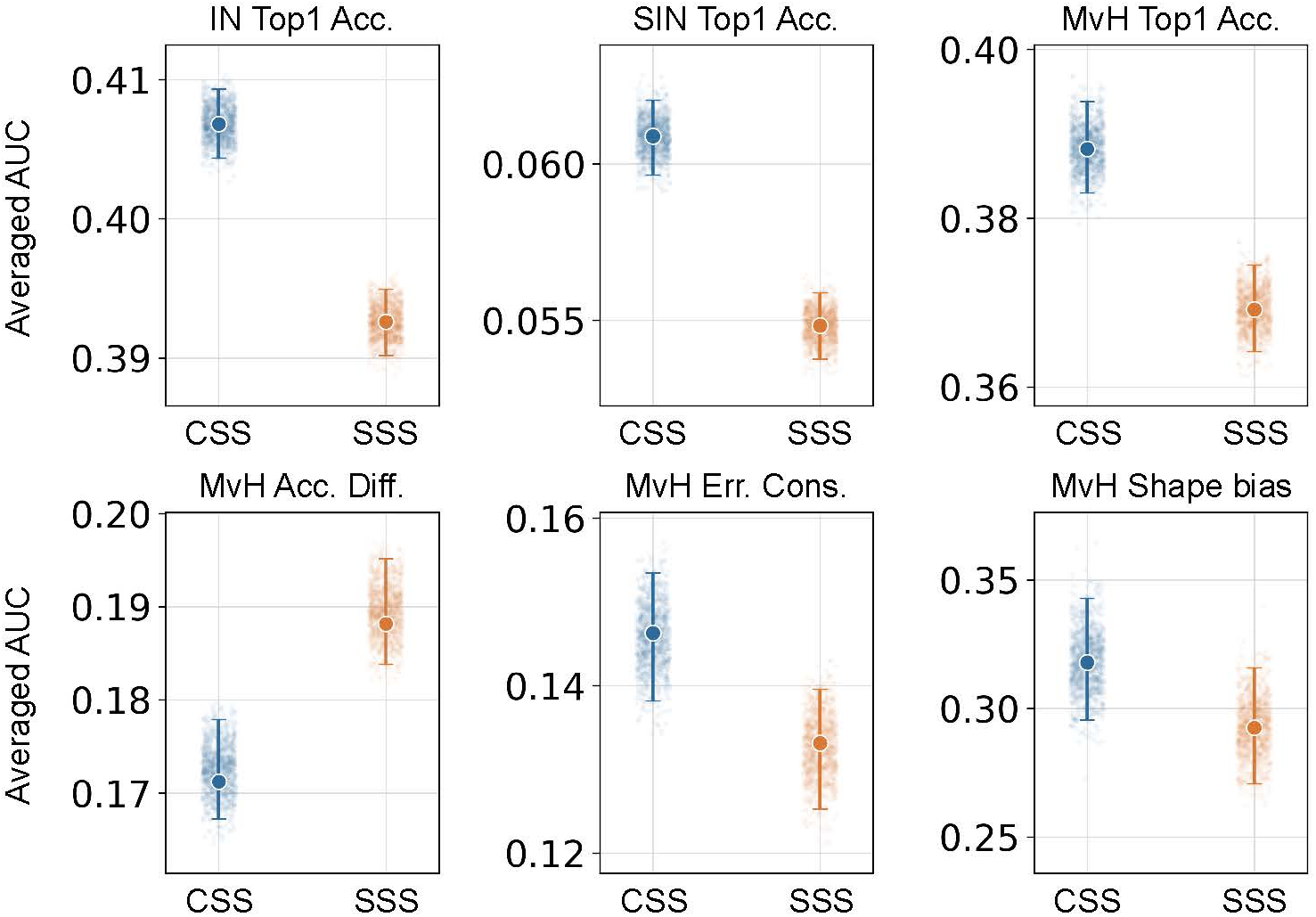
Zoomed-in comparison of CSS- and SSS-based pruning for fine-tuned VGG-19. This figure presents the same six evaluation metrics, aggregation-averaged normalized AUC estimates, and bootstrap distributions as Figure 4, but with y-axis ranges adjusted separately for each metric. The zoomed-in scales make the small differences between CSS- and SSS-based submodels and their 95% bootstrap confidence intervals easier to discern.

### Channel-mask and response-stability analysis of exclusive VGG-19 submodels

We analyzed the channel masks used for the final Gram-matrix-based neural style transfer to quantify how many channels remained after removing channels shared by the CSS- and SSS-based submodels. This analysis supports the matched-pair logic of the behavioral experiment: the final synthesis comparison was intended to isolate features that distinguished the two submodels, rather than features retained by both models. The CSS- and SSS-based submodels were selected from the 30% pruning condition, using mean aggregation for CSS and minimum aggregation for SSS, respectively. The synthesis losses were then computed from exclusive channels after overlap removal. Across the 16 VGG-19 convolutional layers, each selected checkpoint retained 3,869 of 5,504 convolutional channels before overlap removal. Of these retained channels, 3,197 were shared by both checkpoints. Removing the shared channels left 672 CSS-exclusive channels and 672 SSS-exclusive channels, corresponding to an effective retained ratio of 12.2% for each exclusive subspace. The first convolutional layer contributed no exclusive channels because all 64 channels were retained by both checkpoints (Figure 14).

**Figure 14:**
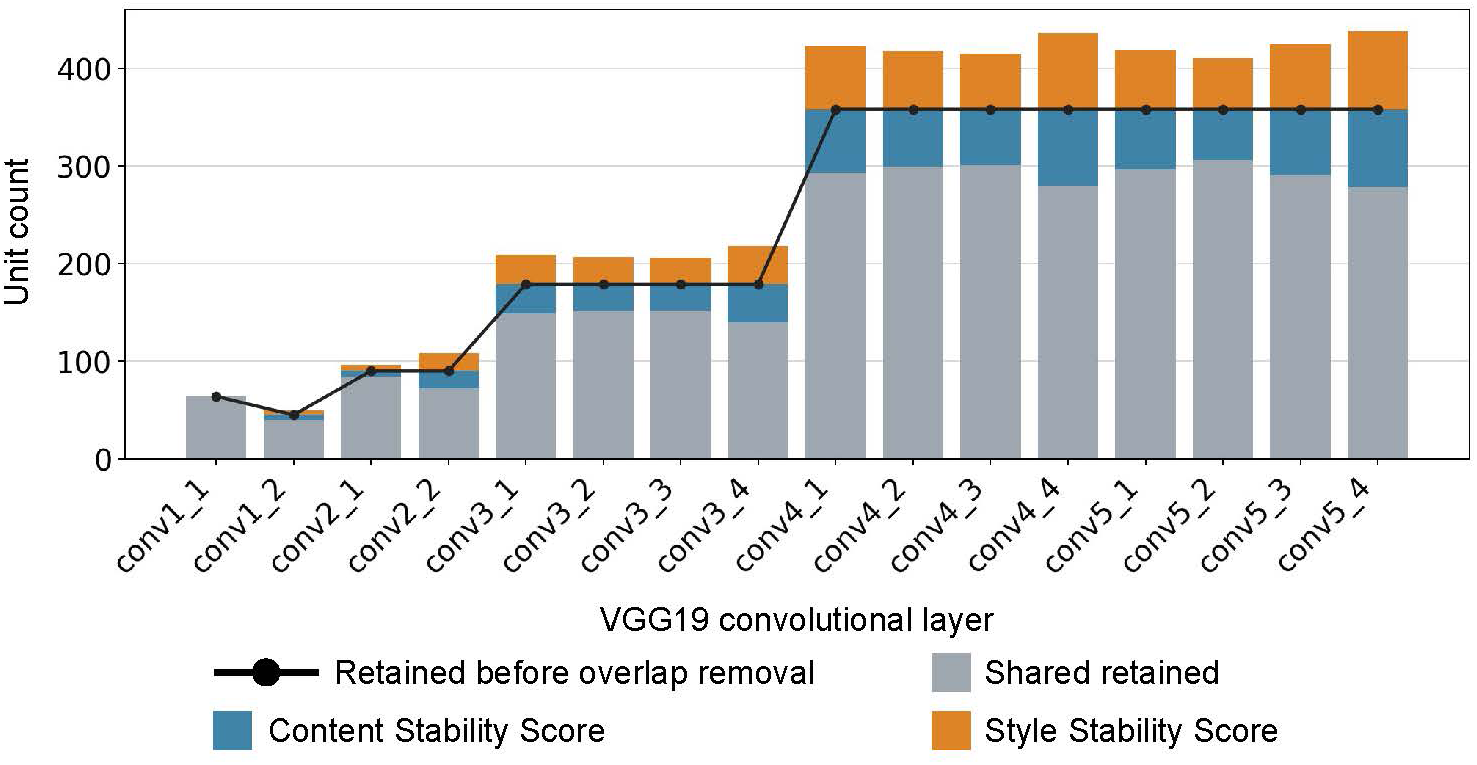
Layer-wise channel-mask analysis for the two VGG-19 based submodels used for Gram-matrix-based neural style transfer. Stacked bars show channels retained by both submodels and channels retained exclusively by either the Content Stability Score (CSS)-or Style Stability Score (SSS)-based submodel after overlap removal. The black line indicates the total number of retained channels before overlap removal. Thus, the gray segments represent channels retained by both submodels, whereas the blue and orange segments represent CSS-exclusive and SSS-exclusive channels, respectively.

To examine the response-stability properties of the channels used for image synthesis, we computed the CSS and SSS of the exclusive channels in the two submodels after fine-tuning. CSS-exclusive channels showed higher CSS than SSS-exclusive channels, whereas SSS-exclusive channels showed higher SSS than CSS-exclusive channels (Figure 15). These differences were consistent across all 15 convolutional layers containing exclusive channels. These results indicate that the exclusive channel sets used for image synthesis exhibited response-stability characteristics aligned with their respective pruning criteria, even after fine-tuning.

**Figure 15:**
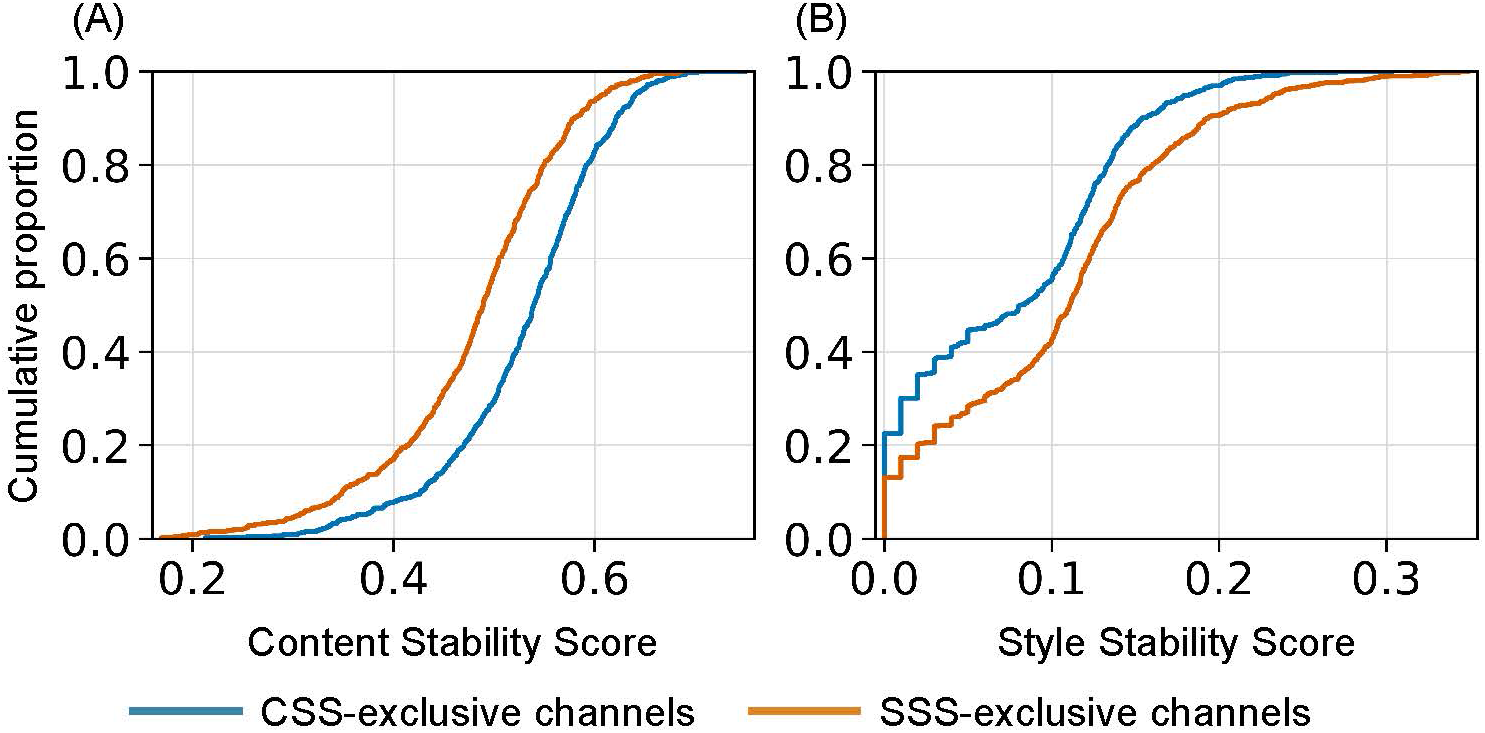
Response-stability distributions of exclusive channels in the fine-tuned VGG-19 submodels. (A) Content Stability Scores (CSS) and (B) Style Stability Scores (SSS) for channels retained exclusively by the CSS-or SSS-based submodel. Each distribution consists of 672 exclusive channels across 15 convolutional layers. The curves show empirical cumulative distribution functions (ECDFs). CSS-exclusive channels showed higher CSS than SSS-exclusive channels, whereas SSS-exclusive channels showed higher SSS than CSS-exclusive channels. The mean differences (CSS-exclusive minus SSS-exclusive) were 0.0511 for CSS (95% bootstrap CI: [0.0504, 0.0517]) and -0.0286 for SSS (95% bootstrap CI: [-0.0302, -0.0155]), based on 200 shared-resample bootstrap replicates over stimulus groups.

### Control analysis of aggregation-rule effects on image synthesis

In the main analysis, the CSS-based and SSS-based submodels were constructed using different aggregation rules: mean aggregation for CSS and minimum aggregation for SSS. Therefore, the observed differences between images synthesized with the two submodels could potentially reflect the choice of aggregation rule rather than the CSS/SSS score itself.

To examine this possibility, we considered four submodel conditions: CSS with mean aggregation (CSS-mean), CSS with minimum aggregation (CSS-min.), SSS with mean aggregation (SSS-mean), and SSS with minimum aggregation (SSS-min.). We then defined two matched comparisons in which the aggregation rule was held constant: CSS-mean vs. SSS-mean and CSS-min. vs. SSS-min. For each comparison, exclusive channel masks were recomputed from the corresponding pair of submodels, and the resulting exclusive feature subspaces were used for image synthesis. The model architecture, pruning rate, fine-tuning stage, input images, style-transfer objective, optimization procedure, and all other synthesis parameters were held fixed across conditions.

Representative examples are shown in Figure 16. This analysis allows CSS- and SSS-based submodels to be compared under the same aggregation rule (CSS-mean vs. SSS-mean and CSS-min. vs. SSS-min.). Across the generated images, no large qualitative differences attributable solely to the choice of aggregation rule were apparent.

**Figure 16:**
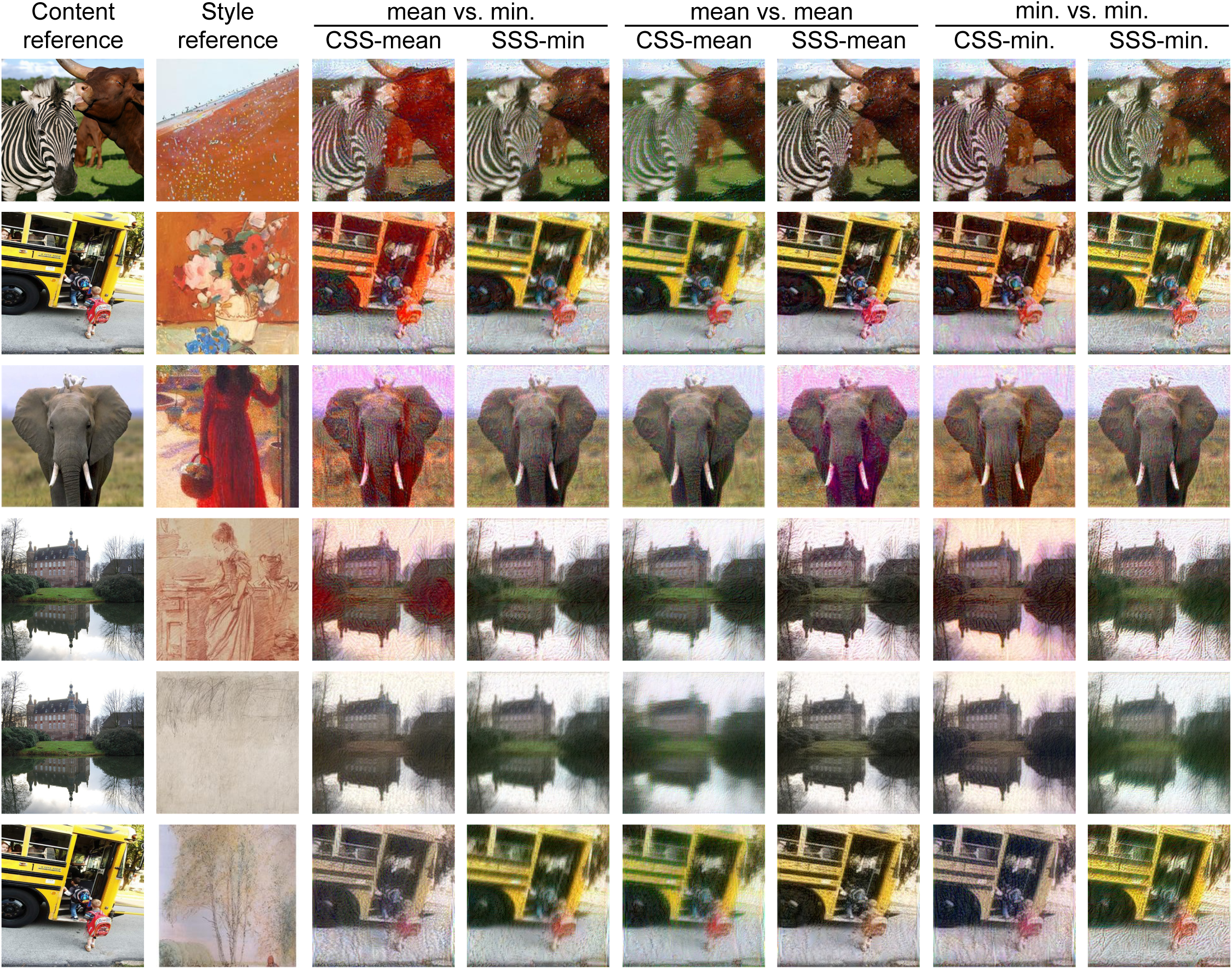
Control analysis of aggregation-rule effects on image synthesis. Representative examples are shown for six content–style pairs also included in the main analysis. For each pair, stylized images were generated using four combinations of submodel criterion and aggregation rule: CSS-mean, CSS-minimum, SSS-mean, and SSS-minimum. Columns show the content image, style image, and three matched pairs of synthesized images: CSS-mean versus SSS-minimum, CSS-mean versus SSS-mean, and CSS-minimum versus SSS-minimum. All synthesis settings other than the submodel criterion and aggregation rule were held fixed across conditions. For comparisons with matched aggregation rules, exclusive channel masks were recomputed separately for the mean–mean and minimum–minimum conditions. All images shown in this figure were either obtained from ImageNet or WikiArt, or generated by applying style transfer to images from these sources.

### Behavioral follow-up analyses

Reaction times were examined as a quality-control check to determine whether the analyzed responses included extremely fast responses unlikely to reflect stimulus-based judgments or unusually slow responses relative to the task. The pre-specified lower and upper reaction time thresholds were 200 ms and 5000 ms, respectively. No analyzed trial fell outside this range; therefore all 5,000 responses were retained for the behavioral analyses (Figure 17).

**Figure 17:**
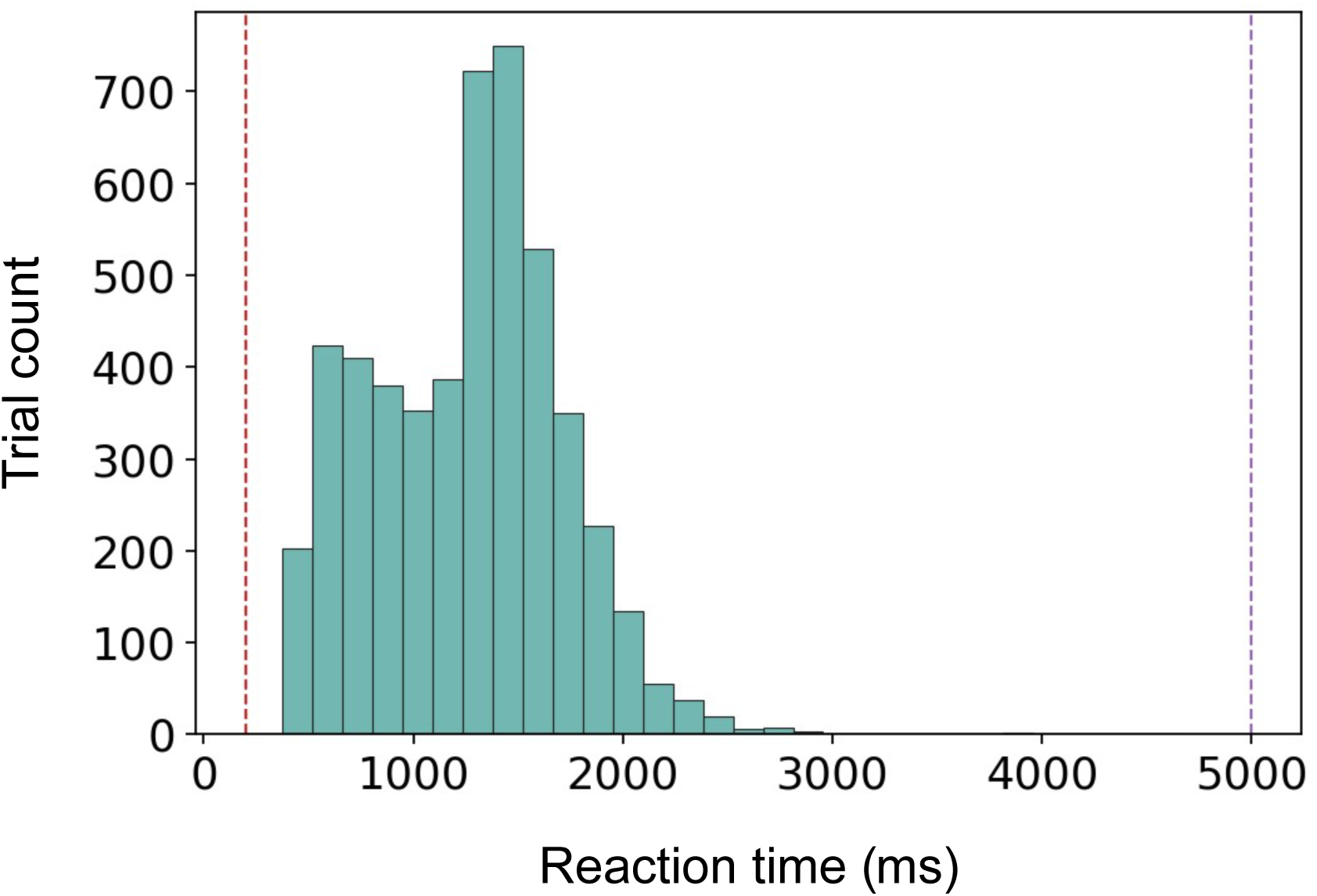
Reaction time distribution for the analyzed behavioral responses. Bars indicate the number of trials in each reaction-time bin. Dashed lines indicate the pre-specified lower and upper reaction-time thresholds of 200 ms and 5000 ms, respectively. No analyzed response fell outside these thresholds.

The main behavioral analysis used a trial-level generalized linear mixed-effects model, complemented by participant-level summaries and content-wise and style-wise analyses. To examine image-specific interactions more directly, we additionally calculated the CSS-based response rate for each content–style combination. This supplementary analysis assessed whether the overall behavioral effect was broadly distributed across the selected stimulus set or was driven by a limited number of image pairs. The resulting matrix revealed substantial heterogeneity across content–style combinations (Figure 18). Several style references yielded consistently high CSS-based response rates across content references, whereas some content–style combinations produced response rates near chance or favored the SSS-based image. These results suggest that behavioral preference was influenced not only by the pruning-defined synthesis subspace but also by the specific combination of content and style references.

**Figure 18:**
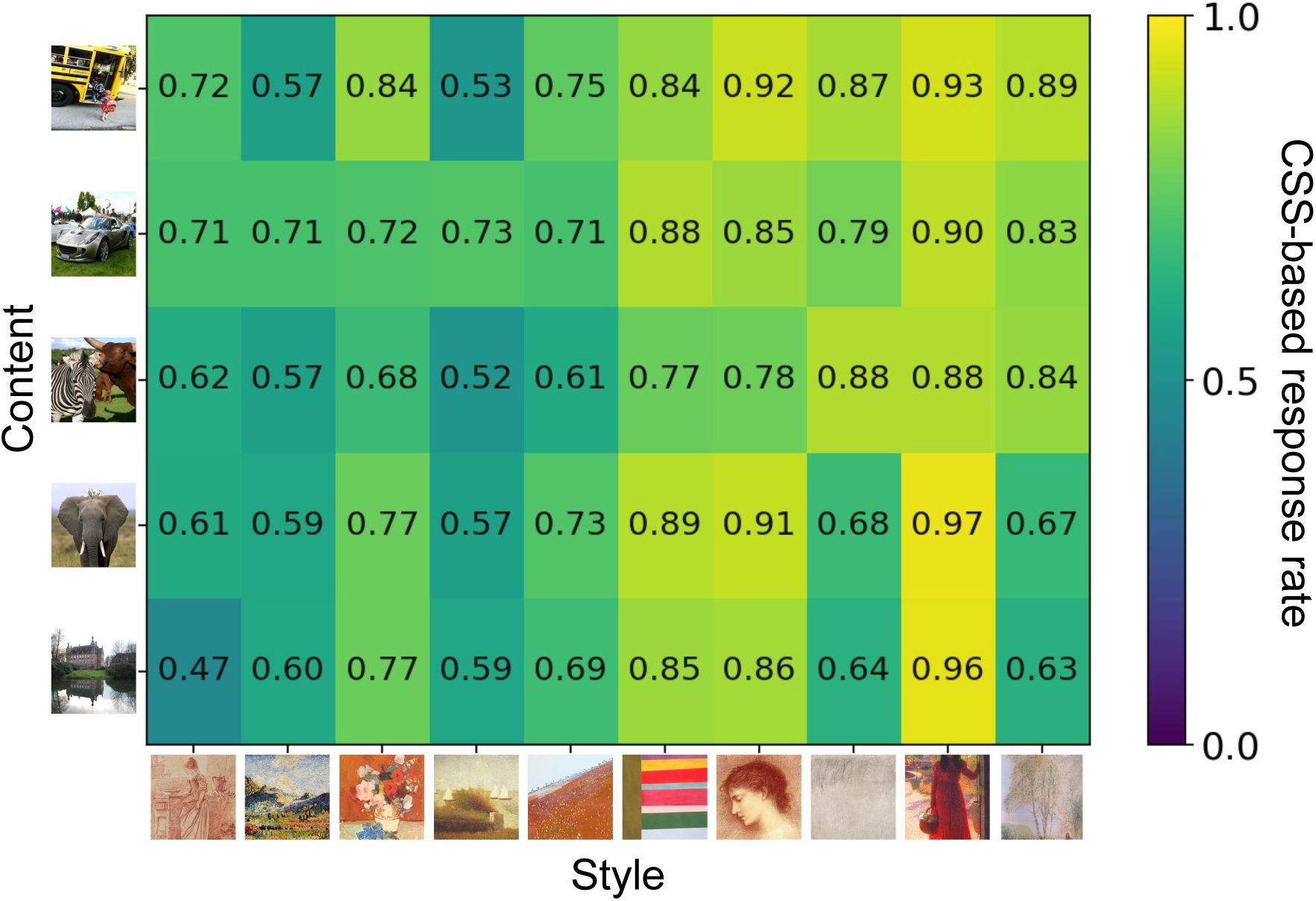
Content Stability Score (CSS)-based response rates for each content–style combination. Rows indicate content references from ImageNet, and columns indicate style references from WikiArt. Each cell shows the proportion of trials in which observers selected the CSS-based image for the corresponding content–style combination. Values above 0.5 indicate a preference for the CSS-based image, whereas values below 0.5 indicate a preference for the Style Stability Score (SSS)-based image. All images shown in this figure were either obtained from ImageNet or WikiArt.

